# Characterising Representations of Hue and Saturation in the Cortex Using Information Decoding

**DOI:** 10.1101/2024.12.19.628842

**Authors:** Ana Rozman, Tushar Chauhan, Jasna Martinovic

## Abstract

One of the enduring questions in the field of colour vision research revolves around how features of colour appearance, such as hue and saturation are represented in the brain. While considerable progress has been made in understanding the transformation of physical colour signals during early processing stages, the mechanisms by which these signals are recombined in later processing, ultimately giving rise to our perceptual experience of colour, remain elusive. A promising avenue for capturing these representations involves decoding from EEG signals. We captured EEG signals in response to 8 evenly spaced isoluminant colours at two saturation levels, taken from a perceptually uniform CIE Lab colour space. Surprisingly, our main finding challenges the expectation that representations of colours that are perceptually more similar elicit more similar signals. Instead, our results show that isoluminant hues perceived as maximally dissimilar tend to evoke the most similar EEG signals, which is in line with predictions derived from opponent theory. Additionally, decoding performs better for lower saturation than for higher saturation hues, contrary to expectations that more distinct hue representations would be associated with highly saturated hues but in line with prediction of magnified differences in latency and amplitude of EEG signals between low-level opponent mechanisms at lower contrast levels. Finally, our findings also highlight the non-uniformity of the cortical representation space for isoluminant colour, indicating that neighbouring hues exhibit varying degrees of similarity. Put together, the results show that low-level features, such as contrast and cone-opponency, drive the EEG response to colour.

## INTRODUCTION

The neural representation of higher-level perceptual properties of colour, such as hue and saturation, remains one of the critical open questions in the literature (Brouwer & Heeger, 2009a). Characteristics of low-level mechanisms in colour vision and response properties of the associated retinogeniculate pathways that represent their neural locus are well-characterised and have been extensively studied (Mollon, 1982; Stockman & Brainard, 2010). We do not yet, however, have a complete understanding of how retinogeniculate signals are further transformed, giving rise to perceptual properties of colour, i.e. hue, saturation and lightness (Conway, 2009; Gegenfurtner, 2003). Whilst perceptual colour spaces characterise colour appearance using these high-level dimensions, they are based on human discrimination data rather than our understanding of neural processes that underpin those discriminations. This is unlike the Derrington, Krauskopf and Lennie (DKL) colour space, which is based on the characterisation of the biological cone-opponent mechanisms (Derrington et al., 1984) but fails to adequately capture attributes such as saturation (Schiller et al., 2017).

Attempts to identify the cortical correlates of opponent hue processing have met with mixed success. Characterisation of early cortical tuning has provided valuable insights, such as the finding that as early as V2, tuning of hue-sensitive neurons becomes much narrower (Kiper et al., 1997). Neurons become progressively more spatially structured in relation to their hue preferences, with nearby regions representing colours of similar hues (Xiao et al., 2003). To a degree, such tuning appears to already manifest in macaque V1 (Xiao et al., 2007). In both macaque and human, hue-sensitive regions in V4 are organised in a circular fashion that mimics that of perceptual colour spaces (Bohon et al., 2016; Brouwer & Heeger, 2009). However, the search for neural correlates of colour-opponent processing has been hampered by confounds due to low-level signal bottlenecks, both in primate recordings (Mollon, 2009; Stoughton & Conway, 2008) and human electroencephalography (EEG; Forder et al., 2017).

EEG research in particular is often limited by a lack of consideration for low-level determinants of the recorded signals. Visual Evoked Potentials (VEPs) derived from EEG recordings are highly influenced by chromatic and luminance contrast and its spatial distribution in the period between 100 and 300 ms (Rabin et al., 1994a; Xing et al., 2015). VEPs reflect V1 cells’ responsiveness to cone-opponency and chromatic contrast, with delayed latencies both for short wavelength-defined stimuli and lower contrast stimuli (Nunez et al., 2017, 2018).

Information decoding represents a more powerful approach to studying the neural representation of hue (Hajonides et al., 2021; Sutterer et al., 2021). Decoding is a multivariate analysis approach, thus allowing for the comparison of multiple relationships between variables simultaneously (Grootswagers et al., 2017). Trials are divided into multiple subsets, with a proportion of these subsets linked to their correct labels and used to train a classifier model. Model performance is then tested on the remaining proportion of trials and evaluated in terms of its success rate in assigning it with the correct label. The more distinct from other stimuli the neural activity elicited by the specific stimulus is, the more likely the success of the model assigning a correct label. The decoding analysis is fitting to the question concerning neural representation of hue. Through representational similarity analysis (RSA), it is possible to evaluate the similarity of the neural distances retrieved from classification with differences derived from other relevant metrics, both key for the question (Ritchie et al., 2019; Ritchie & Carlson, 2016).

Decoding has previously been applied to study the cortical representation of unique and non-unique hues (Chauhan et al., 2023). This study decoded equally perceptually spaced triplets of hues in four regions of colour space: around unique green, yellow, orange and turquoise. For such local neighbourhoods of hues offset by small perceptual differences, decoding was better in unique hue neighbourhoods than in intermediate hue neighbourhoods. This supports the notion of anisotropic cortical hue representation, with larger distances in the vicinity of unique hues. However, this study used a limited number of hues, thus failing to provide a more fine-grained characterisation of the neural code for colour. Likewise, a recent magnetoencephalographic (MEG) study examined the representational organisation of four hues at either positive and negative luminance polarity of equal contrast (Rosenthal et al., 2021), with the same dataset also used to explore the representation of hue and luminance polarity (Hermann et al., 2022). This study reported that both properties can be decoded from MEG data. However, differences in luminance polarity impacted the decoding of hue to a larger extent than differences in hue impacted the decoding of luminance. Luminance signal was found to be more distinct, leading to its overall better decoding performance.

However, differences in information decoding of EEG/MEG signals could also be explained by the aforementioned low-level determinants of VEP waveforms. Adding luminance to chromatic information leads to a pronounced change in the morphology of the VEP waveform (Crognale et al., 2013). Recent research demonstrates hue-specific signals are most successfully decoded when presented at isoluminance, with the hue signatures becoming less distinct from each other with progressive additions of luminance (Chauhan et al., 2023). This is important because luminance does not interact symmetrically with chromatic information from different cone-opponent mechanisms (Martinovic & Andersen, 2018; Shapley et al., 2024). Furthermore, luminance contrast impacts saturation differently for different hues and between the two polarities. Decoding success could thus reflect contrast-driven processes impacting on VEP waveforms, rather than putative high-level colour representations or colour categories.

So far, VEP signals driven by isoluminant hues have only been characterised for relatively small sets of stimuli, looking at limited areas of either cone-opponent or perceptual hue spaces. We aim to expand on this knowledge by comparing signals of hues systematically covering the entirety of the perceptual hue circle. This will allow us to gauge the anisotropy of the neurometric hue space across the entire hue circle. Using changes in chroma for isoluminant stimuli, we also aim to examine how generalisable these cortical hue representations are across saturation levels. In Rosenthal et al. (2021) and Hermann et al. (2022) variations in saturation were not independent of variations in lightness. Here, we compare EEG signals from eight isoluminant hues, equally spaced in perceptual colour space, presented at two saturation levels. Saturation is manipulated through the patches’ chromatic content (chroma). We analyse sets of 2 or 4 hues to compare decoding between hues in their local (neighbouring hues) or global (opposite hues) contexts. We can do this systematically due to the circular nature of the colour space, which we have sampled evenly in our stimulus set, and this allows us to test several predictions. First, we will gain further understanding of the organisation of the neuromeric hue space and its (an)isotropies due to sampling the perceptual hue circle in eight even steps. Second, we will use representational similarity analyses (RSAs), to establish the degree to which cortical hue representations correspond to lower-level (i.e., cone contrast) or higher-level (hue scaling) representational determinants with a higher granularity than previous studies have done. Third, by examining decoding confusion matrices we will be able to further ascertain the determinants of hue-elicited electrophysiological activity: if cortical hue similarities follow the lines of appearance-based mechanisms, proximal hues will be more similar and thus more confusable with each other, but opposite hues being more confusable will point towards opponent-based representation. Finally, if electrophysiological markers of hue representation are encoded independently of contrast, then decoding should be generalisable across levels of saturation. Put together, the study will provide us with a more thorough understanding of cortical signatures of hue and saturation coding.

## METHODS

### Participants

A sample of 15 young participants (4 male) aged between 24 and 37 years (*M* = 27 years) took part in the experiment. All participants had normal or corrected to normal visual acuity, checked with a LogMAR chart for viewing at 63 cm, using a criterion score of 20/20. Participants also had normal colour vision, as verified using the City University Colour Vision Test (Fletcher, 1984). Research procedures were in line with the Declaration of Helsinki, and approved by the University of Aberdeen, School of Psychology ethics committee.

### Apparatus

The experiment was presented on a Display++ monitor (CRS, UK), with participants seated at 70 cm distance. The monitor was positioned in a dark room with no other light source. The experiments were run using the Cambridge Research Systems (CRS) toolbox and CRS colour toolbox (Westland et al., 2012) for Matlab (Mathworks, USA). The presentation was controlled by a VISaGe visual stimulus generator (Cambridge Research Systems, UK) and the display was used in VISaGe mode, ensuring linearised output. Measurements of monitor spectra were obtained using a SpectroCAL (Cambridge Research Systems, UK) spectroradiometer.

### Stimuli

Our stimuli consisted of 2° squares and diamonds (squares rotated by 45°). These were defined in 8 different hues, by specifying Lightness, Chroma and hue (LCh) values derived from a cylindrical transformation of the CIELAB colour space. Colour space conversions were made using the optprop toolbox for MATLAB (*Optprop - a Color Properties Toolbox*, Wagberg, 2024). The LCh colour space was chosen as manipulation of chromatic content in CIELAB provides a reliable measure of saturation, ensuring that colours are equated along this important perceptual dimension (Schiller et al., 2018). Hues were defined by taking cardinal and intermediate directions in CIE LCh space (0°,45°,90°,135°, 180°, 225°, 270° and 315°), corresponding to red, orange, yellow, lime, green, turquoise, blue and purple. Each of the eight hues was presented at two saturation levels, corresponding to chroma levels of 11 (lower saturation) or 22 (higher saturation; see Table 1). The background was set to L=45, equivalent to CIE 1931 xyY coordinates of 0.3127, 0.3290, 14.54 cd/m^2^. Stimuli were set to individual isoluminance using heterochromatic flicker photometry. Cone-contrasts for nominally isoluminant colours as calculated using Smith and Pokorny (1975) cone fundamentals are depicted in Figure 1.

**Figure 1:**
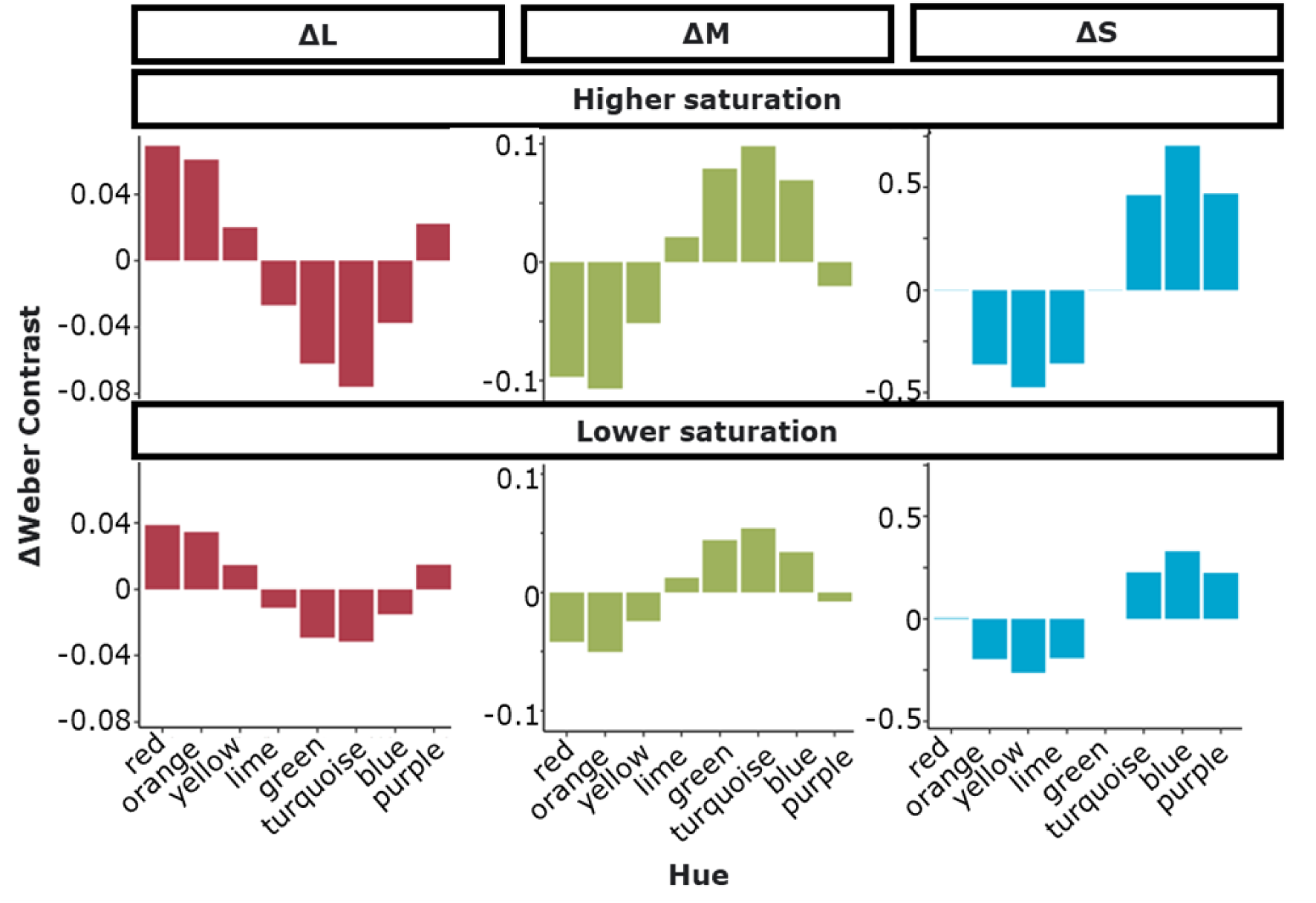
Cone contrasts for Nominally Isoluminant Stimulus colours. Each column shows activations for a specific cone type (L on the left, M in the middle and S on the right), with higher saturation colours in the top row, and lower saturation in the bottom row. Note that the scales differ between cone types but are matched between higher and lower saturation. Cone activations were calculated from spectroradiometric measurements of monitor spectra and Smith and Pokorny (1975) cone fundamentals.

**Table 1.** Stimulus Coordinates in CIE LCh and CIE 1931 Colour Spaces at Nominal Isoluminance.

| L | C | h | x | y | Y |
| --- | --- | --- | --- | --- | --- |
| 45 | 22 | 0 | 0.3667 | 0.3032 | 14.5417 |
| 45 | 11 | 0 | 0.3397 | 0.3161 | 14.5417 |
| 45 | 22 | 45 | 0.4029 | 0.3568 | 14.5417 |
| 45 | 11 | 45 | 0.3575 | 0.3446 | 14.5417 |
| 45 | 22 | 90 | 0.3819 | 0.4018 | 14.5417 |
| 45 | 11 | 90 | 0.3479 | 0.3660 | 14.5417 |
| 45 | 22 | 135 | 0.3211 | 0.4057 | 14.5417 |
| 45 | 11 | 135 | 0.3178 | 0.3659 | 14.5417 |
| 45 | 22 | 180 | 0.2593 | 0.3546 | 14.5417 |
| 45 | 11 | 180 | 0.2859 | 0.3419 | 14.5417 |
| 45 | 22 | 225 | 0.2302 | 0.2908 | 14.5417 |
| 45 | 11 | 225 | 0.2700 | 0.3108 | 14.5417 |
| 45 | 22 | 270 | 0.2453 | 0.2580 | 14.5417 |
| 45 | 11 | 270 | 0.2781 | 0.2926 | 14.5417 |
| 45 | 22 | 315 | 0.2991 | 0.2648 | 14.5417 |
| 45 | 11 | 315 | 0.3064 | 0.2953 | 14.5417 |

*Note.* Actual luminance values were set individually for each observer to reflect their own isoluminance settings.

### Procedure

First, Isoluminance was measured for each observer using heterochromatic flicker photometry (HCFP; Walsh, 1958). Observers were asked to minimise perceived flicker of a 2° foveally presented square stimulus. The adjustments were made using a button box which manipulated the luminance of the flickering colours. Participants completed 8 matches per condition. The highest and the lowest values were excluded and the isoluminance calculated across the remaining 6 values. This value was used to replace the luminance value (Y) in xyY colour space, whilst keeping the hue defining x and y values constant. The matches were acquired for all 8 stimulus colours.

After HCFP, participants took part in an EEG experiment in which they detected a shape oddball. This was either a square or a diamond, counterbalanced across participants. The trial sequence is depicted in Figure 2. Participants were instructed to press a button on the response box as accurately and quickly as possible upon seeing an oddball shape and to withhold their response otherwise. To minimise blink artifacts, participants were asked to blink during the blink cue

**Figure 2.**
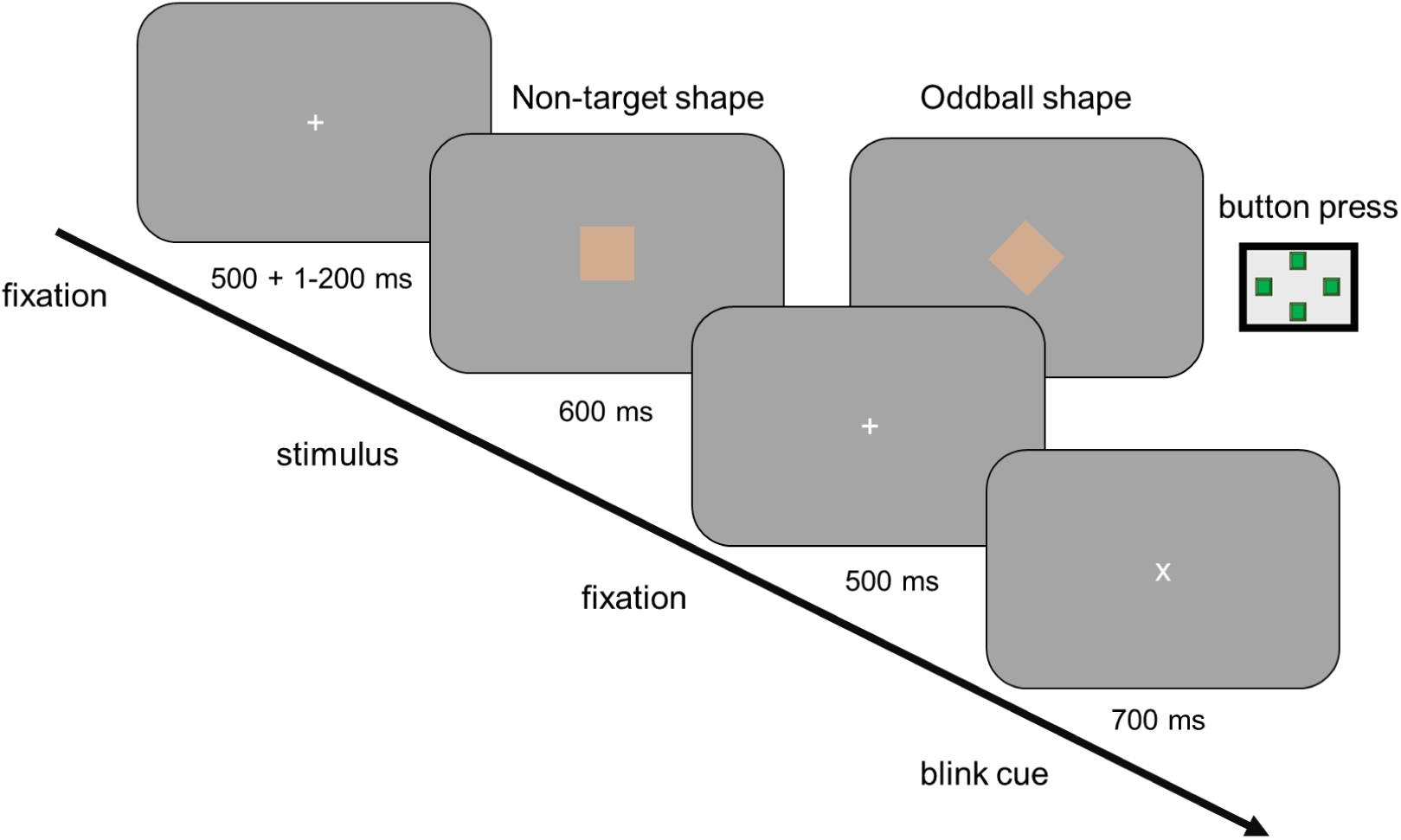
Schematic Representation of a Trial Sequence in the Oddball Task. The trial begins with a fixation cross of randomised duration (500 + 1-200 ms). Subsequently, a non-target or oddball target stimulus is presented (square or diamond, with allocation to target counterbalanced across participants). Participants needed to respond to targets with a button press. Stimulus presentation was followed by a fixation cross, with the responses also accepted during this period (i.e., for up to 1100 ms in total). The trial ended with a blink cue.

(an ’X’ presented at the end of each stimulus sequence) and minimise their blinking and eye movements otherwise. Each hue and chroma level combination was presented 64 times in a randomised order. The oddball shape was present in 15.6% of these presentations (10 oddball shapes, 54 non-target per condition). A total of 1024 trials was divided into 12 blocks (11 with 86 trials and a final block with 78). Each block lasted approximately 5 minutes, adding up to approximately 60 minutes of EEG recording. Participants were encouraged to take breaks between the blocks. Before the recording commenced, a 16-trial practice block was presented to allow participants to familiarise themselves with the task.

After the EEG experiment, we also collected hue scaling data to assess the proximity of stimulus colours to individual unique hue settings. In this task, participants were asked to describe their hue sensation for each stimulus (8 hues at 2 chroma levels) in terms of red, green, blue and yellow components, allocating each of them a score that summed up to 100%. For example, a stimulus matching an individual unique blue setting would receive a 100% scaling for blue sensation and 0% for yellow, green and red, relaying that no other hue sensation is present. On the other hand, a completely balanced intermediate orange hue would receive 50% in red and yellow. A 2° square was presented in the middle of the screen. Each of the 16 stimulus colours (i.e., 8 hues at 2 saturation levels) appeared in randomised order. The task was self-paced, allowing participants to view the stimulus for as long as necessary.

Participants were asked to input proportions of sensation attributed to red, green, blue and yellow separately using a keyboard. All the proportions then appeared on the screen for the participant to review and make any needed adjustments before proceeding to the subsequent trial.

### Electroencephalography (EEG): Data acquisition and processing

Continuous EEG was recorded using a 64-channel BioSemi Active-Two amplifier system, recording the signal at a 1024 Hz sampling rate. 64 Ag-AgCl electrodes were mounted in an elastic, tight-fitting cap. In addition, 4 ocular channels were used to record horizontal (HEOG) and vertical (VEOG) electrooculograms. Data were analysed in MATLAB (Mathworks, US) using the EEGLAB toolbox (Delorme & Makeig, 2004). FASTER (Nolan et al., 2010) and ADJUST (Mognon et al., 2011) toolboxes were used for artifact rejection and correction.

During data pre-processing, recordings were separated into epochs representing individual trials, 1000 ms in duration. This consisted of a 300 ms period before the stimulus onset and 700 ms afterwards. Only the non-oddball trials were included in the data analysis. These presented 864 trials per observer (54 per each of the 16 conditions), or 84.4% of the total 1024 trials recorded for each observer. Furthermore, any trials where observers responded with a button press to a non-oddball trial were excluded. This ranged between 1% and 4%.

Data were filtered using a 40 Hz low-pass and 0.1 Hz high-pass Hamming windowed sinc Finite Impulse Response (FIR) filters, as implemented in EEGlab (Delorme & Makeig, 2004) We also applied a baseline correction between -100 and 0 ms. Contaminated trials were identified and rejected using the FASTER toolbox. This is done by characterising global channel properties using statistical parameters of the data and applying a metric of ±3 *z*-score as a marker for contaminated data. Independent component analysis (ICA) was then performed and the ADJUST toolbox was used to identify specific components driven by blinks, eye movements and local discontinuities. These were removed from the data. Following artifact rejection, FASTER was used to identify any contaminated channels and interpolate them from neighbouring channels. A threshold of ±3 was applied to the *F-*score parameter to define outliers, which were removed and interpolated. A manual check was then performed on the data to verify the quality of artifact rejection. A small proportion of trials was rejected by this procedure, with the remaining non-oddball trials for analysis ranging between 87% and 95% of the total per observer.

To illustrate our grand-mean EEG data, we depict Visual Evoked Potentials (VEPs) and Global Field Power (GFP). This provides an important context for interpreting differences in decoding performance. Previous studies (e.g., Chauhan et al., 2023) have generally found best decoding in the 100 – 300 ms time window following stimulus onset. As discussed in the introduction, this corresponds to the time window where differences in occipital VEPs are most pronounced between stimuli of different contrast (e.g., Nunez et al., 2018). For VEP plots, data were averaged across occipital electrodes, to best capture activity projected from visual cortices (PO7, PO3, POz, PO4, PO6, O1, Oz, O2, Iz).

### EEG: classification

EEG data were analysed using information decoding reliant on a time-windowed error-correcting output codes (tECOC) model (Figure 3). To maximise the data pool, time windows were set up as 20 ms units. This means that at each time point, a 20 ms window of interest was defined, and all data within that window was used to train a classifier at that time-point. The learning units were based on linear discriminant analysis (LDA) classifiers. In human EEG data collected during a similar task, 20ms time-windows have been demonstrated to provide a computationally effective trade-off between accuracy and the size of the parameter space (Chauhan et al., 2023). After the training period, the model was tested by classifying unlabelled data. The process was done for each participant and used signals from all 64 scalp electrodes.

**Figure 3.**
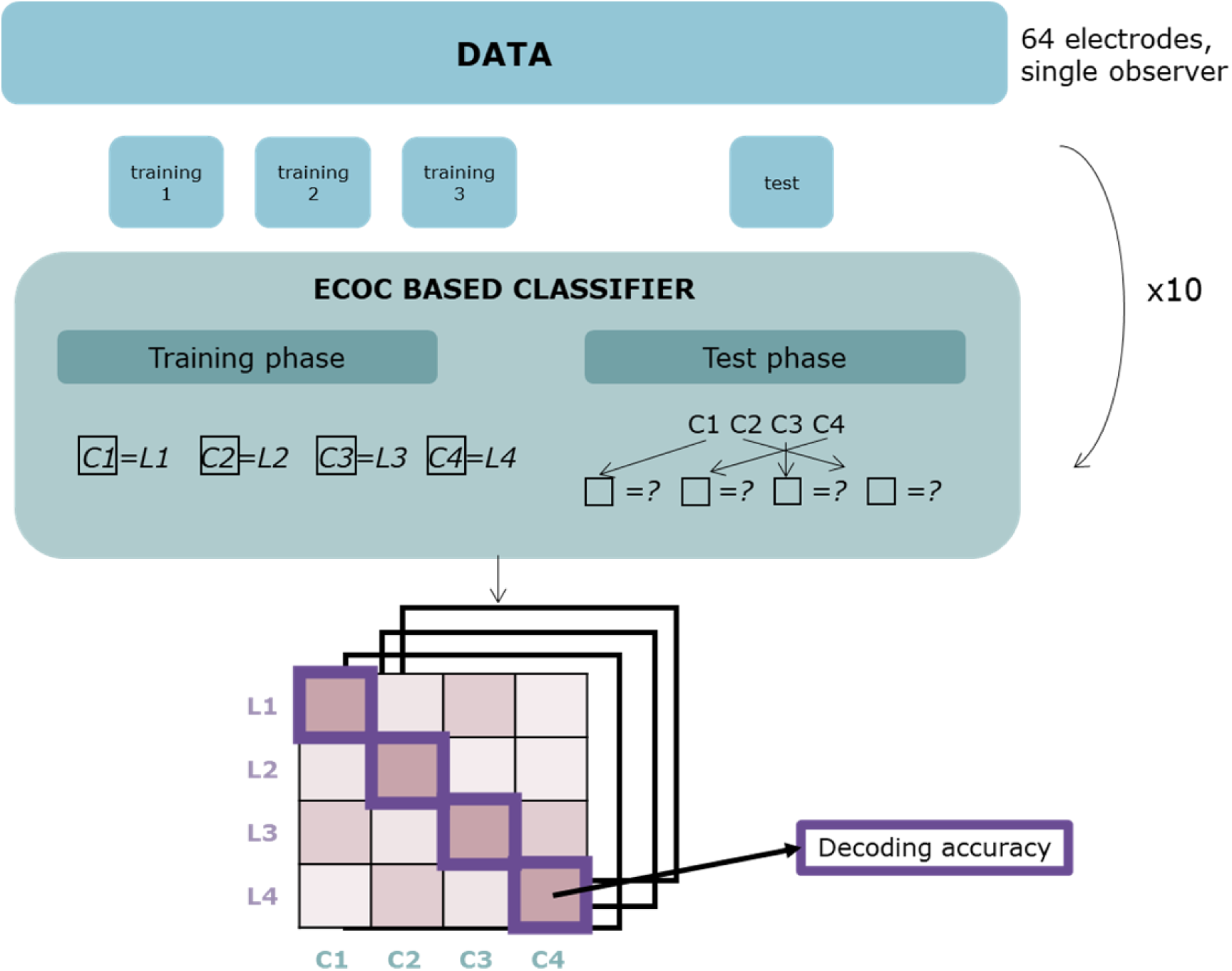
Schematic Representation of the Classification Process for a Single Observer, depicting classification with 4 labels. Data for all 64 channels for each observer is split into 4 folds, 3 of which are used for training. Data from consecutive 20 ms time windows with classification labels are used to train the model. The final fold is assigned classification labels based on training during the test phase. The process is repeated 10 times. The output is a confusion matrix between the actual and classifier-assigned labels across time. The diagonal of this matrix reflects accurate decoding – i.e., when colour 1 is correctly labelled as such – and this is highlighted in the figure using purple squares. The other, incorrectly assigned labels (e.g., labels 2-4 for colour 1) provide information on the degree of confusability between the neural signatures of each colour class.

The data were separated into 4 folds, each containing an approximately equal number of trials. Thus, each trial had an equal chance of allocation into either of the 4 folds. The first 3 folds of data were used to train the model using the stimulus labels. The remaining fourth fold was used for testing the decoder performance. The output of interest was the classification probability confusion matrix generated across the 10 repetitions of the decoding cycle. A permuted baseline of classification probability was created as a control, representing the random chance of the signal being assigned a specific label. For example, in a tECOC model with 4 assigned classification labels, the permuted baseline corresponding to the probability of decoding at each time point should be approximately 25%. For a classification to be above chance, the decoding success of the classifier for a specific label needs to be significantly above this permutated baseline. These differences were evaluated using *t*-tests in the 100- 300 ms window of peak decoding. Multiple comparisons were corrected with the false discovery rate method (Benjamini & Hochberg, 1995).

The confusion matrix represents the proportion of data from each label that is classified into each stimulus category by the model. The decoding accuracy is the diagonal of this matrix (see Figure 3). Thus, the more discriminable the labels are from each other (i.e., the data are less overlapping), the higher the probability that the correct label will be applied in the classification process. On the other hand, labelled signals with more similar characteristics will have a higher probability of being confused with each other, leading to higher error classification rates. The confusion matrix was computed for each time window, thus reflecting decoding success at the specific time point.

### Decoding Data Analysis

As discussed in the Methods section, the outcomes of tECOC signal classifications were the confusion matrices at each time point. We exploited the circularity of the colour space by comparing decoding iteratively in different quadruplet hue sets (verifying the findings on pairwise decoding; see Supplementary Materials). This also limited the complexity of the model so that our dataset (54 trials per label) would be sufficient to train the classifiers. The decoding analysis was performed for four such sets (Figure 4).

1) Distant hues (Figure 4, upper panel): Sets of distant hues allowed us to compare neural signatures of hues with maximal dissimilarity, i.e., those appearing opposite each other in the perceptual hue circle. This analysis was conducted for two sets of four hues. One combined all hues that correspond to cardinal directions of perceptual hue space (red, green, yellow and blue) and another the four hues that appear in between (orange, lime, turquoise and purple). Hues were decoded from distant quadruplets at three saturation levels sets – higher saturation, lower saturation, or combining trials from higher and lower saturation stimuli. To verify that the findings generalise across decoding sets, maximally distant hues were also compared in four sets of opposite hues, e.g., red and green, and lime and purple (see Suppl. Materials).
2) Neighbouring hues (Figure 4, lower panel): Comparing decoding between proximal hues allows us to evaluate the uniformity of the hue space. For example, the representation of a given hue may be more similar to its clockwise than its anti-clockwise neighbour, indicating non-uniformity. Any such departures from uniformity built into the perceptual hue space would be indicative of anisotropic similarity of neural representations along the assumed cortical hue circle.

**Figure 4.**
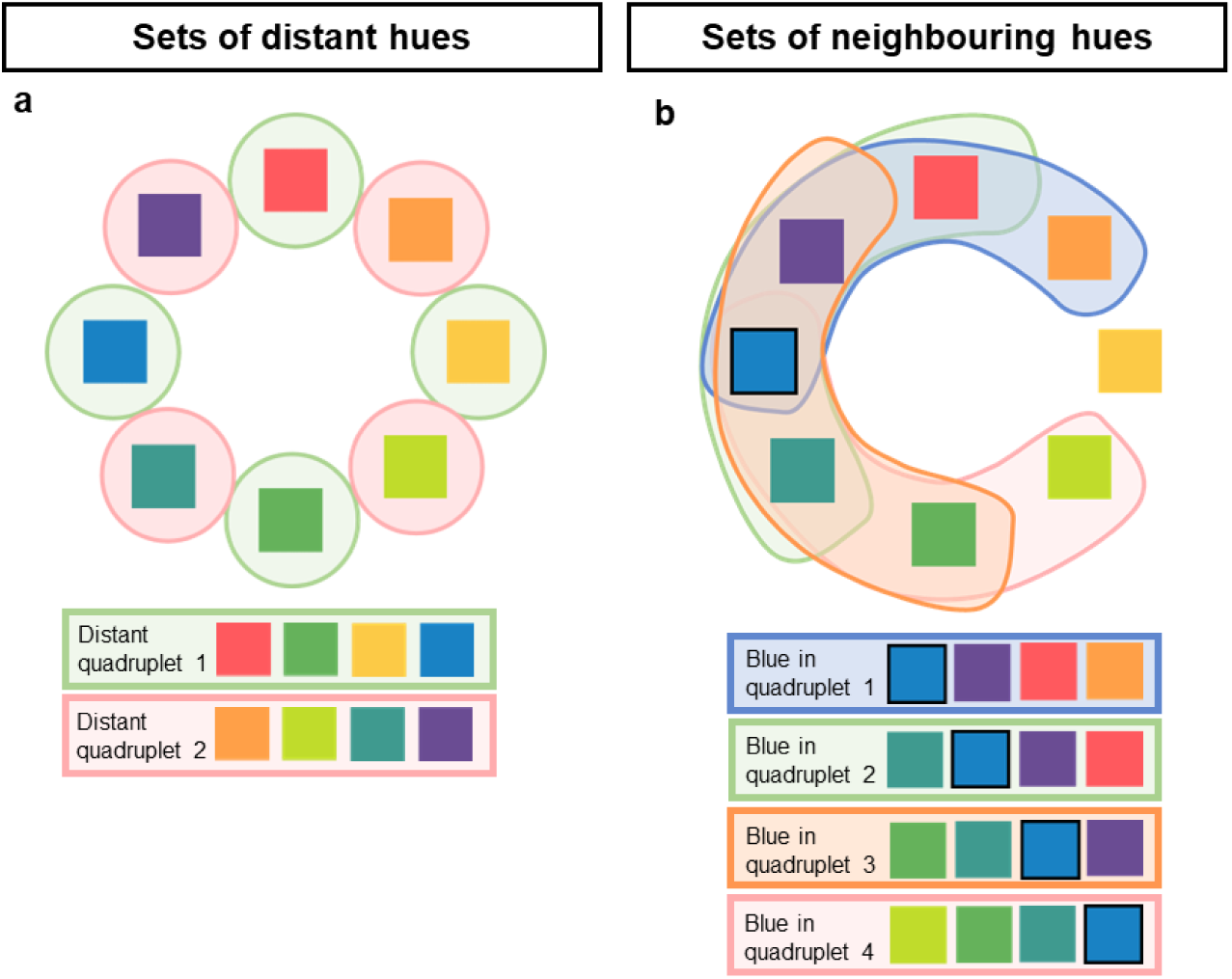
Information Decoding Colour Sets: Distant and neighbouring hues. On the left (a) the sets examining distant relationships between hues are presented. Distant quadruplets are decoded in two sets of hues – first (highlighted in green) combining cardinal hues and second (highlighted in red) the intermediate hue positions. The same distant colours are also examined in decoding pairs, giving rise to a total of 4 comparisons of opposite hues (e.g., red vs. green; see Suppl. Materials). The right panel (b) demonstrates sets where neighbouring hues are compared. This is done (as with sets of opposites) using quadruplets (depicted here) or pairs (Suppl. materials). Each hue is presented in four neighbouring quadruplet sets, once in each of the four conditions. Quadruplets in which the blue hue (outlined) appears are highlighted as an example in panel b. By virtue of the colour space being circular, each colour is flanked by only one other hue in two of the decoding quadruplets, while being positioned more centrally in the other two quadruplets.

**Figure 5.**
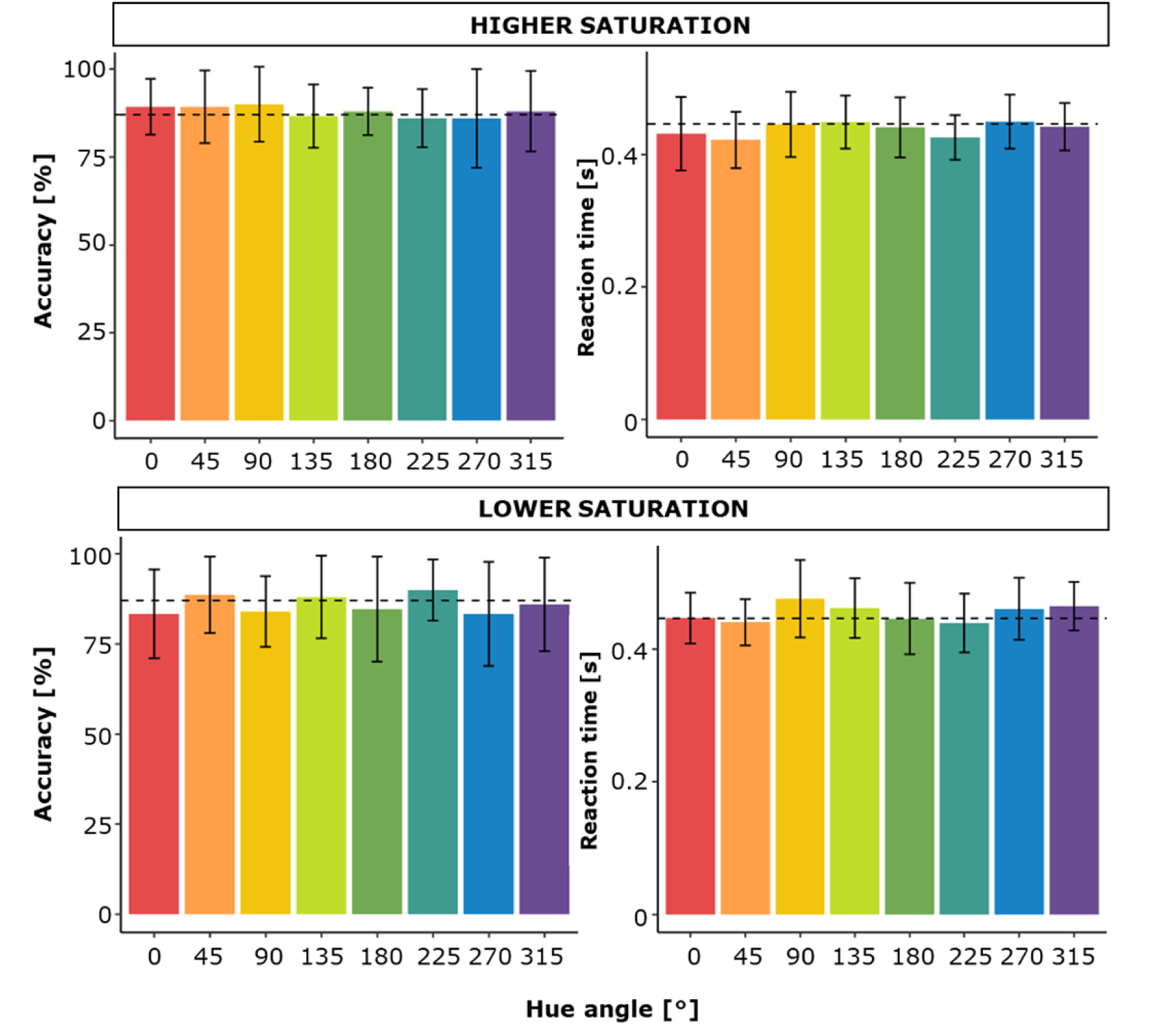
Average Hit rates and Reaction Times (RTs) in Response to Targets. The dashed line denotes the grand average. Note that RTs are faster for more saturated colours, and for orange and turquoise relative to blue, lime and yellow – this is in line with well-known influences of contrast on RTs. Error bars correspond to 2 standard errors.

Neighbourhoods of four hues were defined around the entire circle by moving for a single position each time, e.g., turquoise, blue, purple, red; then blue, purple, red, orange etc. This gave rise to 8 neighbouring quadruplets covering the full perceptual hue circle. Each hue appeared once in each set position, i.e., once as the first, second, third and last item in the set, ensuring that decoding data averaged across colours was not influenced by relative position in the degcoding manifold. Again, decoding was performed for higher and lower saturation level stimuli. For convergent evidence, decoding was also performed on sets of neighbouring pairs (see Suppl. Materials). These were defined using the same approach as neighbouring quadruplets (see Fig. 4)

Statistical analyses were implemented in R (version 4.3.2, R_Core_Team 2023), using packages tidyverse (version 2.0.0, Wickham et al. 2019), lme4 (version 1.1.35.1, Bates et al., 2015), emmeans (version 1.10, Lenth et al., 2019), performance (version 0.12.3, Lüdecke et al. 2021) and DHARMa (version 0.4.6, Hartig 2022). For behavioural data, we fitted binomial data with generalised linear mixed effect models with the fixed effect of hues (red, orange, yellow, lime, green, turquoise, blue and purple) and saturation (low and high) and random by-participant intercepts. Linear mixed effect models with the same structure were fitted to the reaction time data for hits.

For decoding accuracies, a separate linear mixed effect model was fit for each of the two sets: distant quadruplets and neighbouring quadruplets. Decoding accuracy for each hue at a particular saturation level was computed by pooling information from all sets it appeared in. This was done for each individual observer. For example, decoding accuracy for lower saturation orange in neighbouring quadruplets was derived by averaging the accuracy of orange being decoded from orange/yellow/lime/green, red/orange/yellow/lime, purple/red/orange/yellow and blue/purple/red/orange lower saturation level quadruplets. Each model contained fixed effects of hue and saturation, their interaction, and random effect of by-observer intercepts. Simple contrast coding was applied, with arbitrarily chosen reference levels of lower saturation and red hue. Tukey adjustment was used for post-hoc multiple comparisons.

### Representational Similarity Analysis

We also evaluated how the geometry of the classifier results correlated with the representation of our stimuli in a variety of physiologically and perceptually relevant coordinates – cone excitations, relative cone contrasts, opponency mechanisms in DKL space, hue-scaling measurements, and the hue-angle from the LCh space. To do so, we first computed the dissimilarity matrix for the classifier results at each time-point (Chauhan et al., 2023). This was correlated with the dissimilarity matrices of the stimulus in each of the aforementioned coordinates using Kendall’s tau statistic. In both the distant and neighbouring cases the results were averaged over all possible quadruplets – 2 quadruplets in the distant-hues case, and eight quadruplets in the neighbouring-hues case (Fig. 4). This analysis allowed us to compare the geometry of the decoding results over coarse-scale distances (distal-hues) as well as local neighbourhoods (neighbouring-hues).

## RESULTS

### Behavioural data

#### Accuracies and reaction times

In the EEG task, we collected behavioural data including hit rates for targets and target detection reaction times. The response window was limited to 1100 ms and any target trials during which a response was not recorded within this time frame were automatically labelled as misses. We checked if any unreliably fast outliers (<250 ms) were present in the data, but this was not the case.

Binomial data on accuracy for oddball targets (Figure 8, left panel) were analysed using a generalised linear mixed effect model with the fixed effect of saturation level (high or low) and hue and random intercepts for participants. None of the fixed effects contributed to the model (saturation: χ2(1) = 1.984, p = .159; hue χ2(7) = 3.082, p = .877; interaction χ2(7) = 5.555, p = .593; for full model details, see Supplementary materials).

Reaction times for hits (Figure 8, right panel) were analysed using a linear mixed effect model with saturation level (high or low) and hue as fixed effects and random intercepts for participants. The two fixed effects contributed additively to the model (saturation: χ2(1) = 19.717, p < .001; hue χ2(7) = 32.962, p < .001; interaction χ2(7) = 4.762, p = .689; for full model details, see Supplementary materials). Post-hoc tests revealed faster RTs for more saturated colours, and for orange and turquoise relative to blue, lime and yellow hues (orange vs. yellow t(233)=-4.09,p=.001,vs. lime t(233)=-3.327,p=.023, vs. blue t(233)=-3.299,p=.025; turquoise vs. yellow t(233)=-3.905,p=.003; vs. lime t(233)=3.137,p=0.040, vs. blue t(233)=-3.109, p=.044). This is in line with established RT differences for isoluminant colours (Mckeefry et al., 2003), with more contrast leading to speeded responses, and a general slowing of RTs for colours more aligned with the S-(L+M) axis as opposed to the L-M axis.

#### Hue scaling

All cardinal hues (red 0°, green 180°, blue 270° and yellow 90°) were, on average, rated as having more of their respective cardinal content (Figure 9). This exceeded 75% for red, yellow and blue. Green was scaled as having more green (61%, with 95% CIs of 53% – 68%) but this was closely followed by blue (33%, with 95% CIs of 14% - 54%).

Inspection of Figure 6 makes it clear that hue scaling is very similar at the two chroma levels. This is confirmed by statistical analyses, which show that the only determinant of scaling is hue (see Supplementary Materials for full statistical tables). Focusing on the asymmetries in appearance of yellowness, redness, blueness and greenness across CIE LAB, a few differences are notable: 1) blueness is most broadly distributed, with both turquoise and blue having a high and statistically not robustly different blueness (90% for blue, with a 95% CI of turquoise’s difference to blue being -11% to 7%); 2) CIE LAB green has high blueness content, so that the maximum scaling for greenness is given to lime (61% greenness for green, with lime being higher by 10% – 29% (95%CI); 3) yellowness is the most ‘peaky’ and the least variable between individuals (effect size being nearly identical with and without random effects), with orange and lime both receiving below 50% yellowness ratings; 4) red is comparably less ‘peaky’ and is also more variable between individuals (conditional r^2^ = 0.48, marginal r^2^=0.32), with both purple and orange having substantial perceived redness.

**Figure 6.**
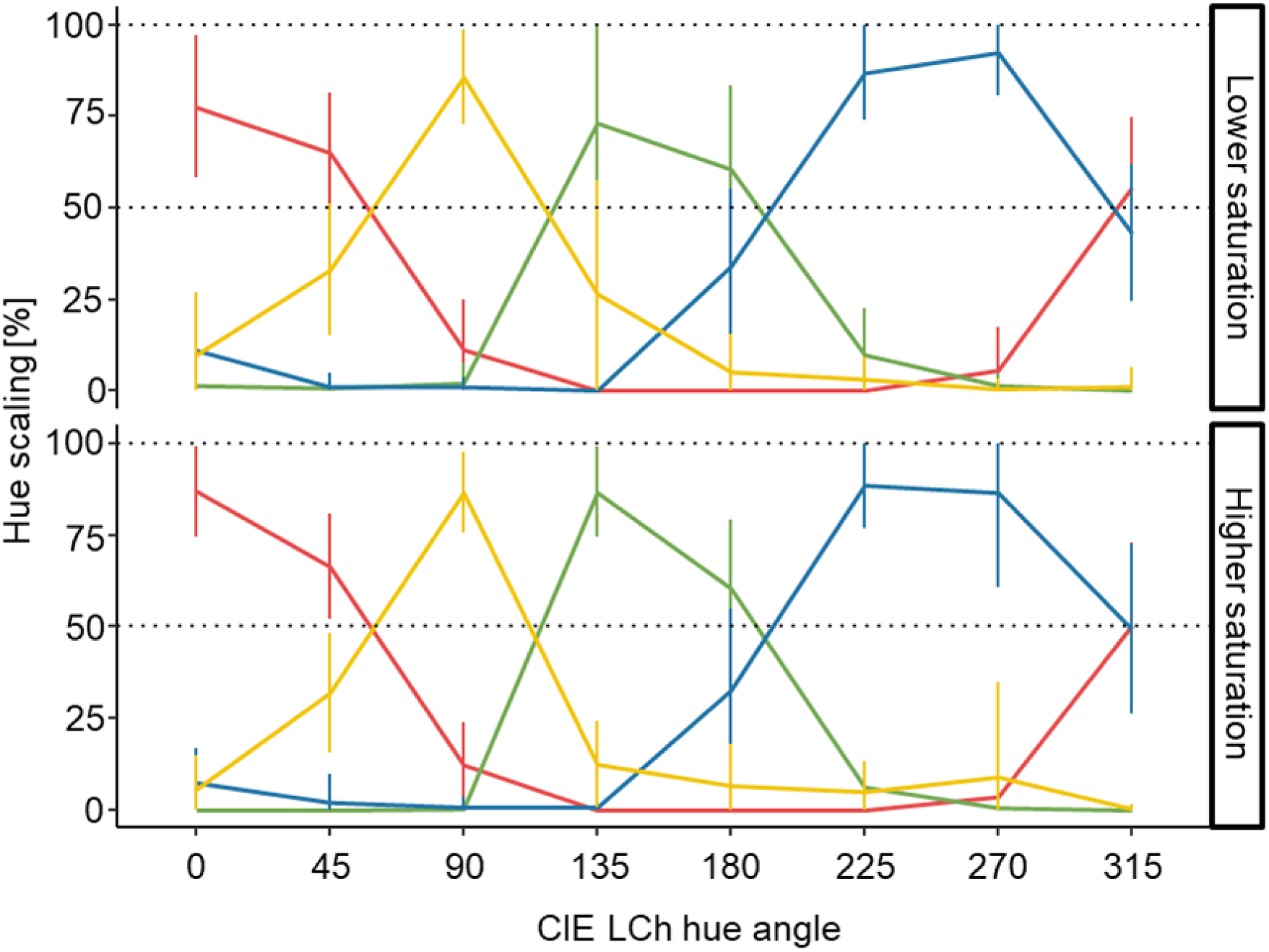
Mean Hue Scaling results. The lines show average percentage ratings for each of the four reference hues (red, green, blue and yellow). Top row shows hue scaling for lower and bottom row for higher saturation. Error bars represent 1 standard deviation and are capped at 0 and 100%. Dashed lines correspond to 100%, which would be an idealised unique hue sensation, and 50%, corresponding to an idealised mix of two proximal hues. Multiple departures from these ‘ideal’ scaling levels confirm that whilst CIELAB may be uniform in how it represents discriminability, its axes do not correspond to cardinal axes of colour-opponent (i.e., unique hue) mechanisms.

**Figure 7.**
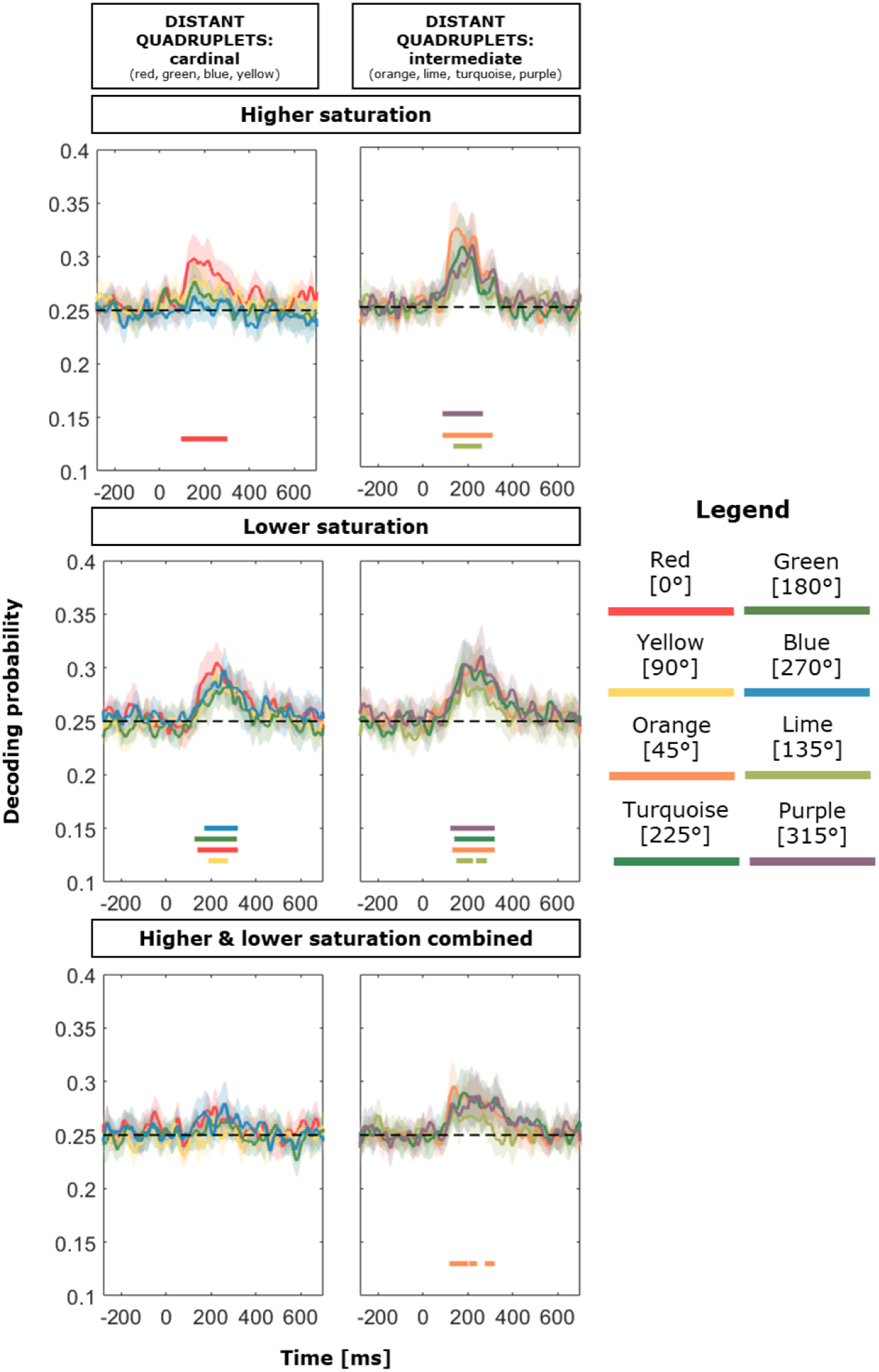
Decoding Accuracy for Distant Quadruplets. Decoding accuracies are presented for three saturation conditions (higher, lower, or combined higher and lower saturation). The outcome is the accurate decoding probability across time (i.e., classifying a stimulus as red when it was indeed red, etc.). Each hue’s decoding probability was compared to a permuted baseline during the 100 – 300 ms window. A horizontal line indicates the time course of significant differences at the bottom of each plot, presented in the same colour as the hue. Error envelopes show 95% confidence intervals.

### EEG decoding: Distant sets

We compared the decoding performance of each hue to its permuted baseline (Figure 10; chance level 25%) within each of the sets (cardinal or intermediate) and saturation level (higher, lower, or combined across saturation).

When hues were presented at higher saturation, red was the only colour that was decoded significantly better than the permuted baseline between 100 and 300 ms following stimulus onset. Whilst we also observed a slight uptick in decoding probability for green and yellow during the same time window, this was not significantly different from the baseline. At higher saturation, decoding was better for the set containing non-cardinal distant hues - purple, orange and lime were all decoded successfully. For the remaining hue, turquoise, a small decoding peak was observed in the same time frame, but this was not statistically significant.

Decoding performance become better at lower saturation. We reliably decoded all four unique and intermediate hues. The decoding peak again occurred in the 100-300 ms time window. However, the hue signals became less distinct when two saturations were combined: whilst a slight uptick in decoding probability was observed for non-cardinal distant hues, only orange decoding was significantly better than baseline. This indicates that cortical signals associated with each label are likely to be distinctive at each saturation level (see VEP section below).

More information about the decoding performance can be gauged by examining the incorrect labels the model assigns rather than by just focusing on the correct labelling. We present these off-diagonal elements from the confusion matrices in Figure 8. Decoding probability is listed for each colour from the decoding sets at each saturation level. The label that has the highest probability of being assigned is the one the signal is most characteristic of. For example, if the model has comparable probability of attributing two hue labels to a certain hue signal, that signal is characteristic of both labels to a similar extent.

**Figure 8.**
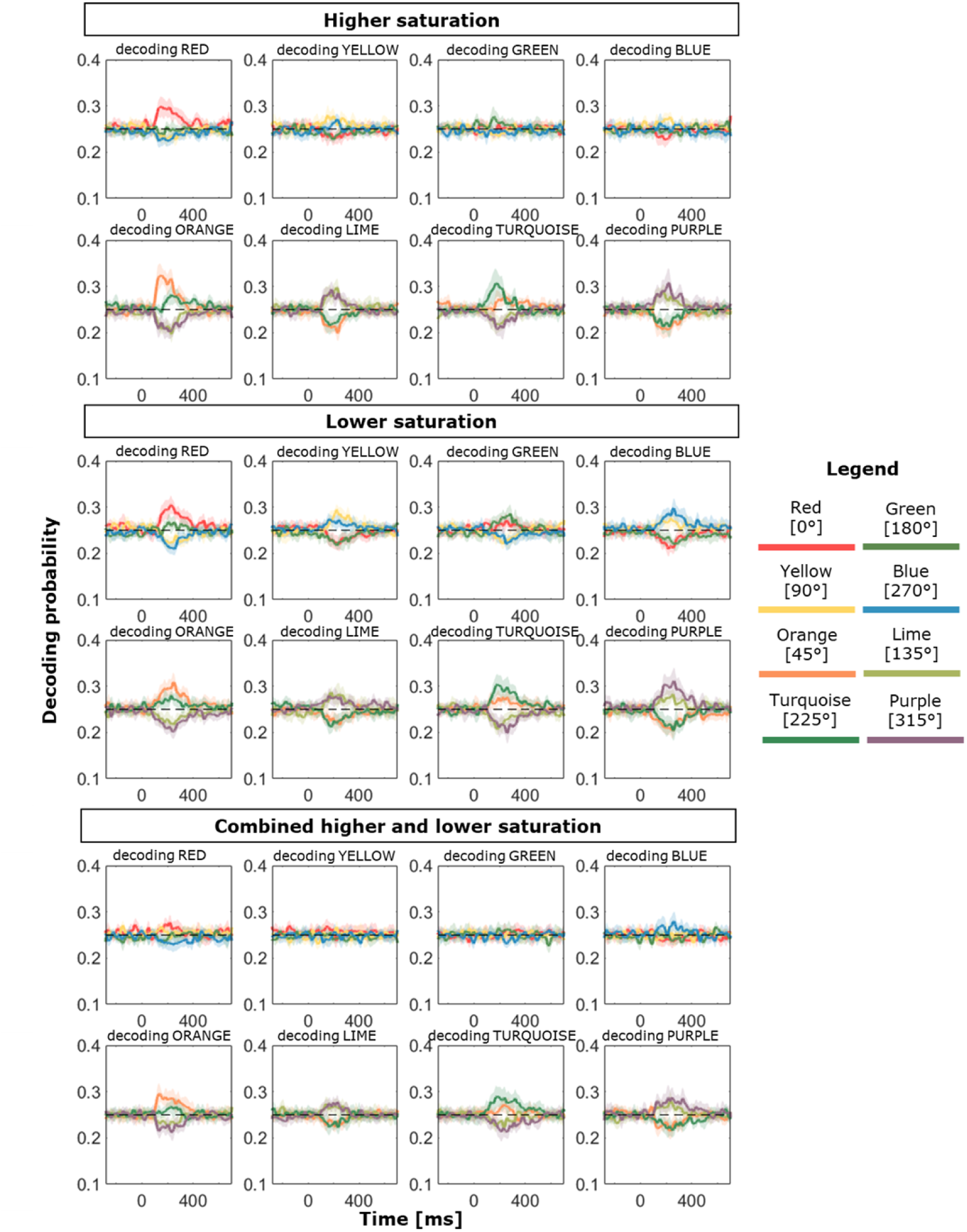
Full classification results for Distant Quadruplets. Confusion matrices provide information about signals that are more confusable with each other. The panels show the probability of any label being assigned to a signal across time for each decoded colour class. A dashed line denotes the chance level (0.25). A separate plot for each class is presented at each saturation level. Accurate decoding (e.g., red label assigned to red class) is the same as that depicted in Figure 7, but this figure casts further light on representational similarity. In cases of successful decoding, the correct label is more similar to (i.e., more confusable with) the opposing hue (e.g., red is more confusable with green and less confusable with blue and yellow). Error envelopes show 95% confidence intervals.

At high saturation condition, red was decoded from the other hues in the set, which were all at chance level. This differed from the set of distant hues at higher saturation, where the correct hue label was the most commonly attributed one in each condition. Interestingly, for all four non-cardinal hues, the correct label was also closely followed by its opposing label. Purple was most frequently misclassified as lime and vice versa, and turquoise was most frequently misclassified as orange and vice versa. This was mirrored by a reduction in labelling probability for the other two hues That means signals of opposing hues are most similar to each other, and the signals of the two proximal colours the most dissimilar.

Highest decoding probability overall was observed at lower saturation. Again, the correct hue was most frequently labelled in each condition, followed by the opposing hue within the same axis. For cardinal hues with combined saturation levels, labelling performance remained around chance level for all hues. The confusion pattern was again replicated, with the correct hue label most frequently attributed, closely followed by the opposite hue.

Decoding accuracy (i.e., the data along the diagonal of the confusion matrix) in the 100-300 ms window for distant quadruplets was compared between all hues at the two saturation levels using a linear mixed effects model (for full details of the best fitting model, see Supplementary Materials). Saturation and hue interacted in predicting accurate decoding (χ2(7)=15.232, p=.033; Figure 9).

**Figure 9.**
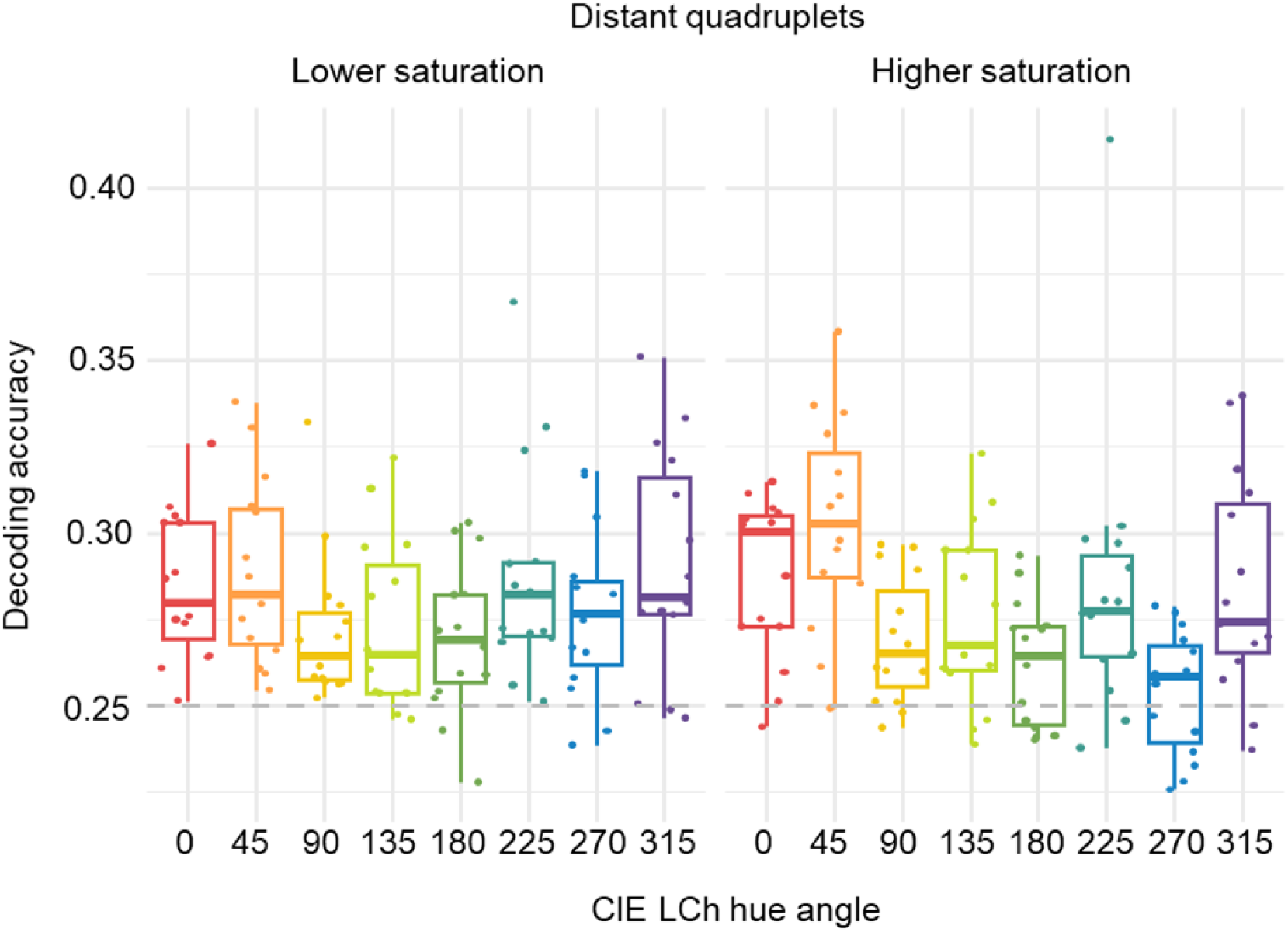
Box plot of accurate decoding proportions over the 100-300 ms window for distant quadruplets. Horizontal lines are medians, b*oxes encompass 50% of the distribution (the interquartile range – IQR), the minimum and maximum whisker values extend the box by -/+ 1.5 * IQR to depict variability in each direction, while dots show individual participants*. Horizontal dashed line at 0.25 denotes the chance decoding level. Note that decoding is (1) highest for high saturation orange, likely driven by higher confusability of orange and turquoise at lower saturation (see Fig. 8) and (2) lowest for high saturation blue, which cannot be decoded above chance level (Fig.7), likely due to its high similarity with yellow (Fig. 8).

While orange at the high saturation level was most decodable, blue at the high saturation level was the least decodable. While colour decodability mostly follows the warm/cool hue split, with red, orange and purple amongst the highest decodable and green, blue and lime amongst the least well decodable, there are two notable exceptions: yellow is grouped with the less well decodable colours, while turquoise belongs with the better decodable colours (see post-hoc tests in Supplementary Materials).

We also performed decoding of distant (i.e., opposite) pairs, but it did not produce significantly distinguishable outcomes (for more details, see Supplementary Materials). This additional finding should be considered in the context of the colour confusion lines presented above. Hues in quadruplets benefited from the presence of two lateral hues from which their signal could be decoded, while signatures for opposite pairs turned out to be too similar in isolation.

### EEG decoding: Neighbouring sets

To complement the characterisation of neurometric hue space derived from distant quadruplets, we also compared representation in neighbourhoods of perceptually proximal quadruplets and pairs (see supplementary materials). In each case, these were established at each level of saturation by moving around the perceptual hue circle in steps of one, defining eight quadruplets and eight pairs for each hue (see Fig. 4).

Similar to the decoding of distant quadruplets, higher saturation proximal hue quadruplets had lower decoding success than lower saturation quadruplets.

Within high saturation quadruplets (Figure 10), sets combining blue, purple, red and orange, and red, orange, yellow, and lime were most distinct, with all four hues successfully decoded from the baseline in the 100-300 ms window. These were followed by purple, red, orange and yellow; turquoise, blue, purple and red; and orange, yellow, lime and green quadruplets, each with three hues decoded successfully. Only purple was successfully decoded from the green, turquoise, blue and purple group. In the two ‘green region’ quadruplets between yellow and blue (yellow, lime, green and turquoise; and lime, green, turquoise and blue), only lime was decoded successfully.

**Figure 10.**
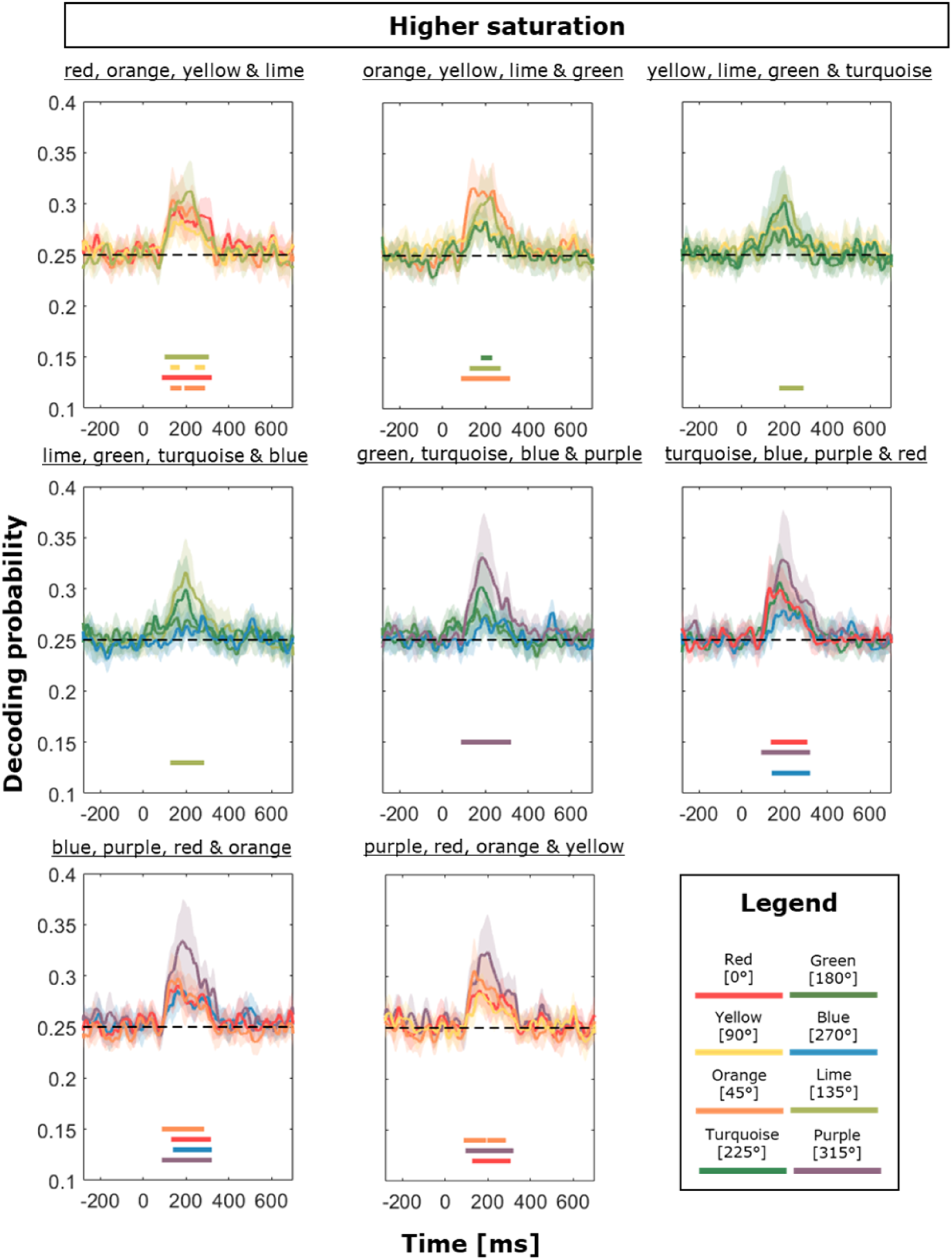
Probability of accurate Decoding for Neighbouring Quadruplets at Higher Saturation. Lines in the 100-300 ms window show significant differences against a permuted baseline. Decoding was closest to chance for the quadruplets containing lime, green and turquoise. Error envelopes show 95% confidence intervals.

At lower saturation (Figure 11), significant decoding of all four hues in a quadruplet was possible for all but the three quadruplets extending into the ‘cool’ region (i.e., green/blue hues) of the colour space (orange, yellow, lime and green; yellow, lime, green and turquoise; and lime, green, turquoise and blue). Within these quadruplets, the colour that could not be decoded was changing.

**Figure 11.**
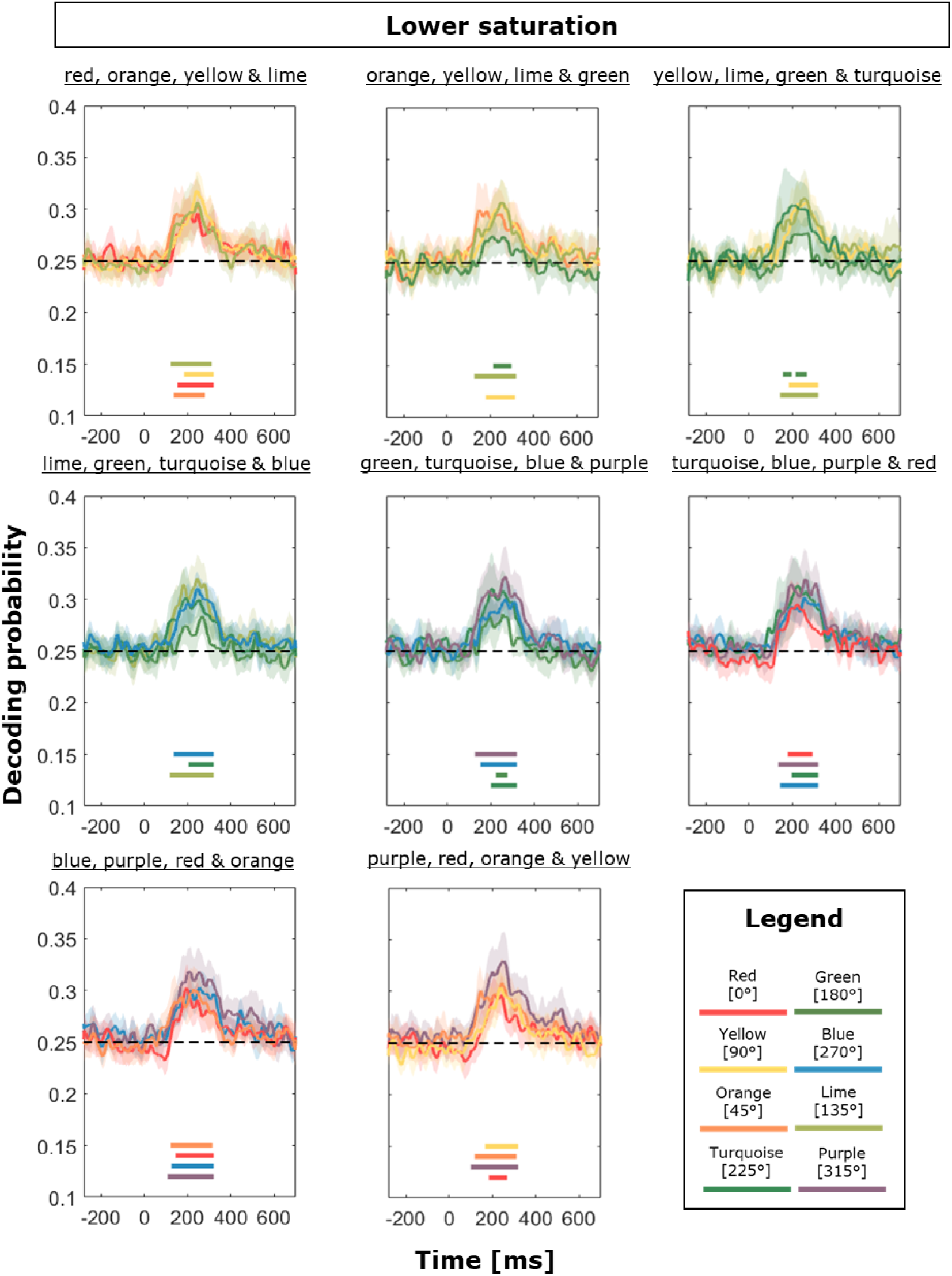
Probability of accurate Decoding for Neighbouring Quadruplets at Lower Saturation. Lines in the 100-300 ms window show significant differences against a permuted baseline. Comparisons with Figure 10 reveal that decoding was better for less saturated hues, although the patterns still differ between different colour space regions, releaving underlying neurometric asymmetries. Error envelopes show 95% confidence intervals.

For example, the green hue could be successfully decoded when in a group with orange, yellow and lime (i.e., as the outer element in the quadruplet, only neighbouring lime), and could also be decoded above chance when in the neighbourhood of yellow, lime and turquoise, but it was no longer decodable when it moved into a higher blue content neighbourhood with yellow replaced by blue (i.e., lime, green, turquoise and blue).

Examination of confusion patterns reveals further details as to the origin of these local asymmetries in representation. At higher saturation, the highest peak for labelling almost always corresponded to the correct label (Figure 12). The exception to this was the green hue which was misclassified to a similar degree as to which it was successfully classified with at least one of its neighbours in any of the four quadruplets it appeared in. Most mislabelled were the orange and red when they appeared in the same quadruplet. But some hues were less confusable with their neighbours - overall, the highest labelling success at high saturation was achieved for purple. This means the signal associated with the purple label is less easily confused by that associated with any of its neighbouring hues, regardless of its position within a set.

**Figure 12.**
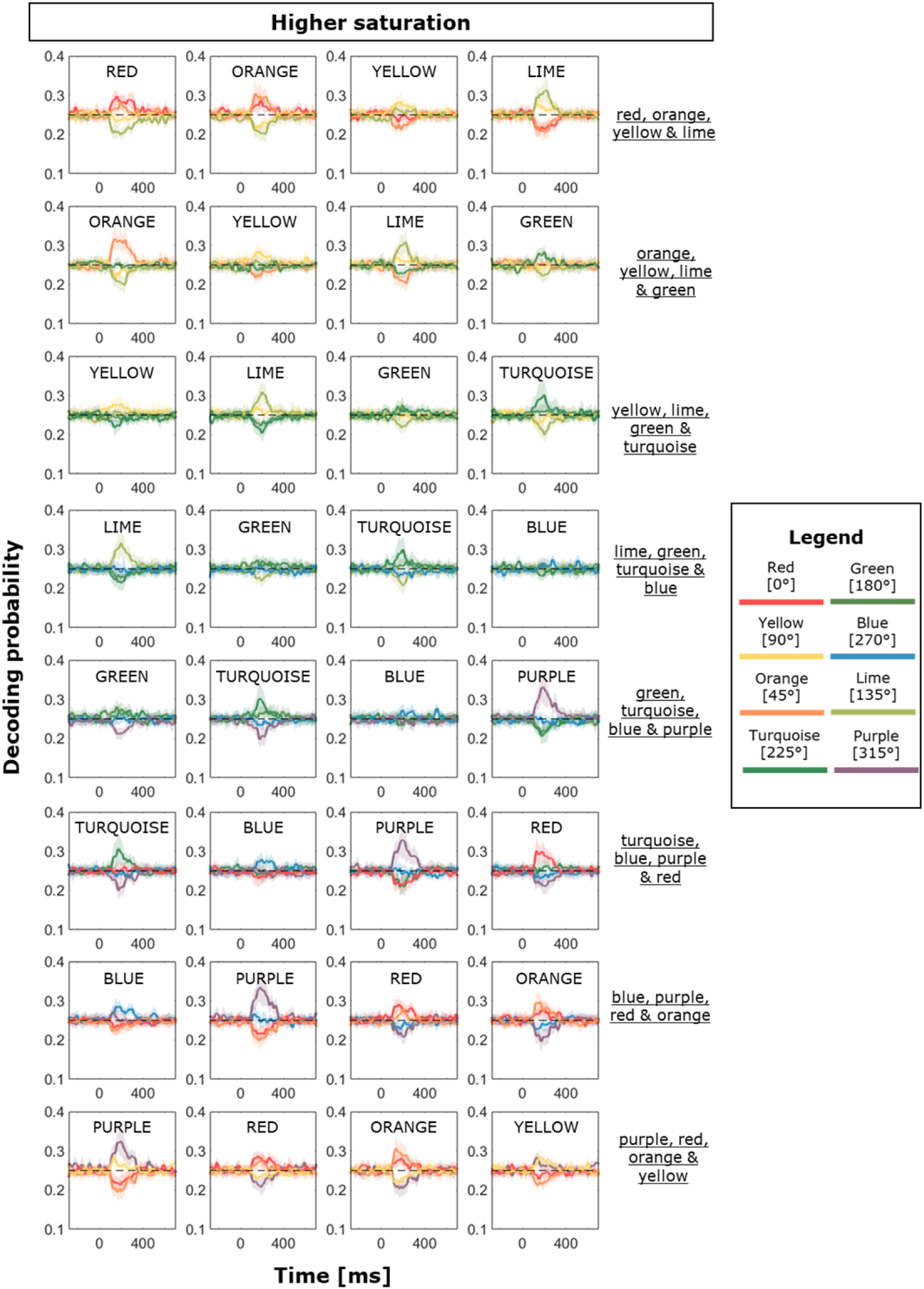
Full classification results for Higher Saturation Neighbouring Quadruplets. The correct label of the hue is shown in each subplot. The panels show the probability of any label being assigned to a signal across time for each decoded colour class, thereby visualising the full content of the confusion matrices. A dashed line denotes the chance level (0.25). Accurate decoding (e.g., red label assigned to red class) is the same as that depicted in Figure 10, but this figure casts further light on representational similarity. Compared to decoding from distant hue sets (Figure 8), colours appear to be less frequently mislabelled in their neighbourhood quadruplets. A prominent exception is the red hue, frequently mislabelled as orange and vice versa. Error envelopes show 95% confidence intervals.

At lower saturation, the green hue was frequently mislabelled as turquoise in groups where the two appeared together (Figure 13). Peak labelling success was higher for blue at lower compared to the higher saturation - blue was not commonly mislabelled as any other hue within comparison quadruplets. As was the case at higher saturation, the most considerable confusion between labelling hues remained between red and orange, with purple again consistently achieving the highest labelling probability peaks among all colours.

**Figure 13.**
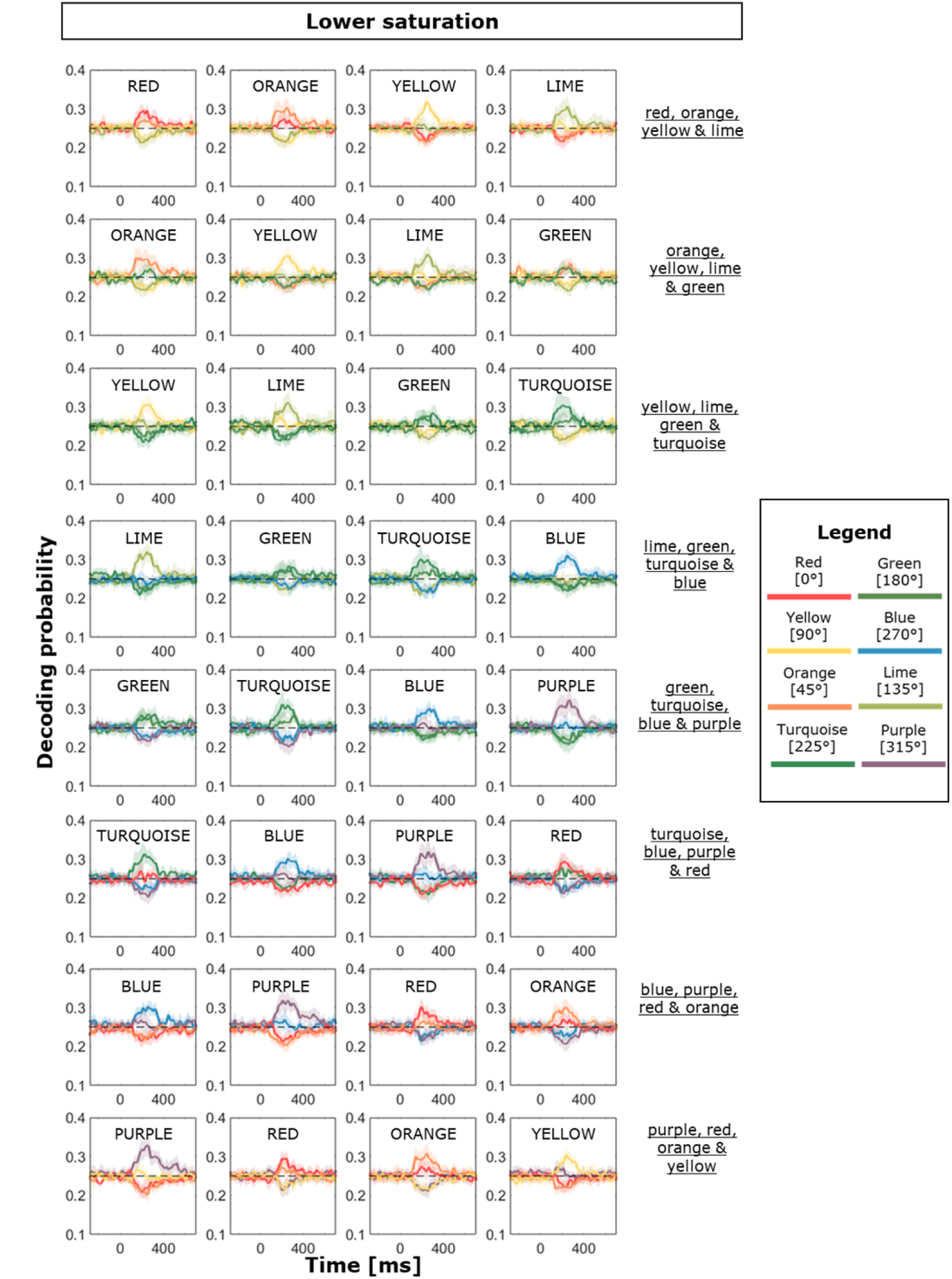
Full classification results for Lower Saturation Neighbouring Quadruplets. The correct label of the hue being decoded is shown in each subplot. The panels show the probability of any label being assigned to a signal across time for each decoded colour class, thereby visualising the full content of the confusion matrices. A dashed line denotes the chance level (0.25). Accurate decoding (e.g., red label assigned to red class) is the same as that depicted in Figure 11, but this figure casts further light on representational similarity.

Similar confusions noted for higher saturation colours persist (e.g., red and orange). Higher peaks in labelling probability reflect the higher overall decoding performance at lower saturation, compared to higher saturation (depicted in Fig. 12). Error envelopes show 95% confidence intervals.

We analysed decoding probabilities for all neighbouring hues at the two saturation levels using a mixed effects linear model (for full details of the best fitting model, see Supplementary Materials). On average, decoding was reliably above chance (95%CI: 27.5 – 29.1%, with 25% being chance level). Again, we found an interaction between saturation and hue – decoding accuracy was substantially different between low and high chromatic contrast (χ2(7)=36.3, p<.001; see Fig. 14) across different colours, with blue undergoing a statistically significant decrease when moving from low to high contrast decoding quadruplets. Decoding for green was poorer across both levels of saturation, while purple was associated with somewhat higher decoding outcomes overall (for post-hoc tests, see Supplementary Materials).

**Figure 14.**
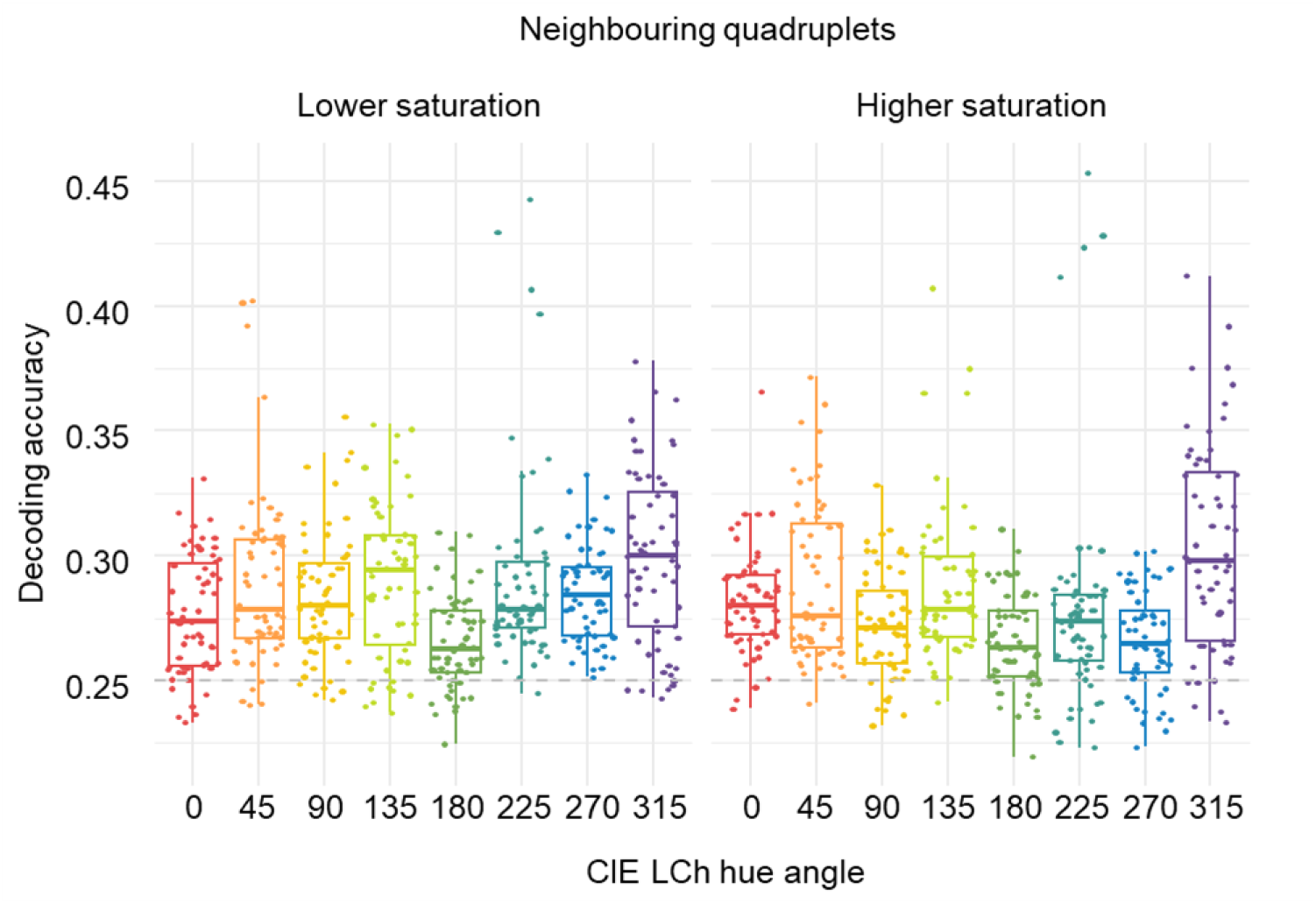
Accurate decoding proportions over the 100-300 ms window for neighbouring quadruplets. *Boxes encompass 50% around the mean, while dots depict individual participants*. Note the (1) drop in decoding for more saturated blue, (2) overall poorer decoding for green and (3) overall better decoding for purple. Horizontal dashed line at 0.25 denotes the chance level.

### EEG analysis: Visual Evoked Potentials

Differences between signals on which decoding is based can be better understood by comparing their VEPs at occipital sites (see Supplementary materials for a demonstration of correlations of decoding outcome with occipital difference waves). An overview of VEPs reveals differences between low and high saturation colours, in line with the expected effects of chromatic contrast on EEG waveforms (e.g., higher early amplitudes for colours with more L-cone content; Knoblauch et al., 1998; see Figure 16). The VEPs also highlight differences between hues in latencies and magnitudes during the time window of the contrast-driven P1/N1 complex. Inspection of the GFP plots – a useful indicator of signal-to-noise ratio (Koenig & Melie-Garcia, 2010) - reveals two areas of significant signal increase, corresponding to P1 and N1 peaks, which coincide with the time window of highest decoding success, i.e., between ∼100- 300 ms.

We also performed representational similarity analyses (RSAs) with various low-level and high-level determinants of colour contrast and appearance, but neither of these properties alone could sufficiently explain the underlying neural representational space obtained in our EEG study (see Supplementary Materials). This is unsurprising, as hue scaling does not depend on contrast, but our waveforms do; similarly, if perceptual distances explained the decoding outcomes, then we wouldn’t have obtained highest confusability lines along cone-opponent lines.

## Discussion

We investigated the neurometric hue space using information decoding from EEG data collected in response to the passive viewing of isoluminant hues. We successfully decoded hue representations from sets of perceptually distant orthogonal hues and sets of perceptually more similar proximal hues. The decoding success was better at lower than higher saturation levels and revealed marked non-uniformities in the neurometric space. When signals were compared for hues positioned opposite each other in perceptual hue space, decoding was possible for groups of four (i.e., combining two opposite poles) but not for two opposite hues themselves. We also find further asymmetries in the neurometric space, with between-hue representational differences compressed in the green region of the colour space. These findings provide a first detailed characterisation of cortical hue representation as measured by EEG, encompassing the complete perceptual hue circle sampled at 45° intervals. The findings demonstrate that the neurometric hue space is non-uniform and that decoding of hue is largely driven by contrast and opponency. Put together, our findings point towards considerable contributions from low-level, contrast-driven factors that need to be taken into account when interpreting the decoding of cortical hue representations using EEG data.

Changes in chromatic content will elicit changes in VEP waveforms in terms of their latency and amplitude (Nunez et al., 2018) and changes to these VEP attributes can drive decoding (Martinovic, 2022). As shown in our VEP and GFP plots, stimuli with lower contrast produce a delayed latency of the first peak which is also sustained for longer than for higher contrast stimuli (Nunez et al., 2018), in agreement with previous accounts in the literature (Crognale et al., 1993; Rabin et al., 1994b). With higher contrast, these VEP responses become more similar to each other, which is likely the reason why hue-related signals are more distinguishable at lower compared to higher contrast. After all, information decoding is just a more powerful way to analyse EEG data by reparametrizing it into an information measure, harnessing the data from multiple electrodes to place the decision criterion at a point that allows most accurate labelling. With linear discriminant analysis, which is often used to allow for more interpretable decoding outcomes, this corresponds directly to a point at which the difference waves are maximally different (for a demonstration of correlations between simple difference waves and information decoding using our data, see Suppl. Materials).

On the contrary, a high level explanation of the observed decoding patterns would need to refer to perceptual or categorical qualities of the stimuli used by our study. Since highly saturated colours are more commonly chosen as better prototypes of their category (Olkkonen et al., 2010; Witzel, 2019), this might lead to the expectation that decoding would to be better at a higher saturation level which is the opposite of what we observe. Whilst the effect of categorical representativeness on decoding was not directly examined by our study, the unique hue content ratings collected from our observers did not suggest substantial differences between the two saturation levels. Despite this, neural activity significantly differs between them. In addition to the evidence of the strong influence luminance contrast has on decoding success (Chauhan et al., 2023), our findings emphasise the importance of chromatic contrast over and above any putative higher-level (e.g., colour-opponent) representations of colour.

A comparison of neural signals associated with perceptually distant hues, located opposite each other in the perceptual hue space, revealed a strong influence of opponency on cortical hue representations. For sets of four hues, each hue is decodable against its orthogonal hues in the set, whereas labelling confusion is driven by its opposing hue in the set. This is confirmed further, as decoding pairs of opposite hues is impossible. These analyses imply that signals of opposite hues are more similar to each other than those of proximal hues. Neuroimaging studies of primates suggest many colour-sensitive cells in V1 are tuned to linear colour opponency, inheriting it from cone-opponent signals from LGN, before neurones start exhibiting narrower tuning in extrastriate cortical areas (Engel et al., 1997; Gouras, 1974; Kiper et al., 1997). Our findings of opponency could reflect some of this processing, as the signatures would be most similar for bipolar axes in terms of their cone-opponent activation. Whilst not fully aligned with cone-opponent processing axes, opposing pairs in CIE LAB space still oherit to an extent a degree of cone-opponency, as evident from their cone excitations (see Figure 5.1). Throughout the study, decoding success was best in the time window between approximately 100 and 300 ms following stimulus onset, with a peak roughly at 200 ms. This time course coincides with the marked influence of V1 single- and double-opponent cell activity on chromatic VEP waveforms (Nunez et al., 2018). Thus, our decoding success can be directly linked to changes in VEP forms, resulting from activation of V1 neurons tuned to cone-opponency.

In addition to opposite hues, our stimuli also allowed for comparing decoding between proximal hues. This has revealed non-uniformities in the neurometric hue space. The colour sets encompassing the same range of perceptual dissimilarity (as reflected by the CIE LAB distance) vary in decoding success depending on the section of the perceptual hue circle they occupy. For neighbouring sets with four hues, decoding was least successful in neighbourhoods containing green hues and those where orange and red appeared together. Decoding within neighbouring pairs confirmed this, with all neighbouring perceptual hues being successfully decodable, except for the green and turquoise and the red and orange (see Supplementary Materials). The chromatic VEPs for L-cones have higher contrast gain than those for M-cones (Knoblauch et al., 1998). In line with this, the VEP waveforms for our orange and red stimuli are characterised by a higher P1 than other hues (see Fig. 15) - and cone activations for our study’s red and orange hues demonstrate that these are the stimuli with the highest L content (see Fig. 1). Therefore, the inability to successfully decode two signals characterised by a similar chromatic VEP driven by L-cone activation from each other would suggest that decoding is based on distinguishing between these properties. Furthermore, the two other neighbouring hues with the most similar L-cone activation to each other are the green and turquoise hues, which also have neural signatures that are not decodable from each other. An alternative explanation could be that these stimuli appear perceptually more similar to observers than stimuli in neighbouring pairs within which decoding is successful. However, the hue scaling scores of our observers reveal that green and turquoise are more dissimilar in terms of their unique hue content than turquoise and blue. The latter pair is, however, decodable with higher success. Low-level processing influences on chromatic VEP waveforms remain the most parsimonious explanation for the asymmetries observed in the neurometric hue space.

**Figure 15.**
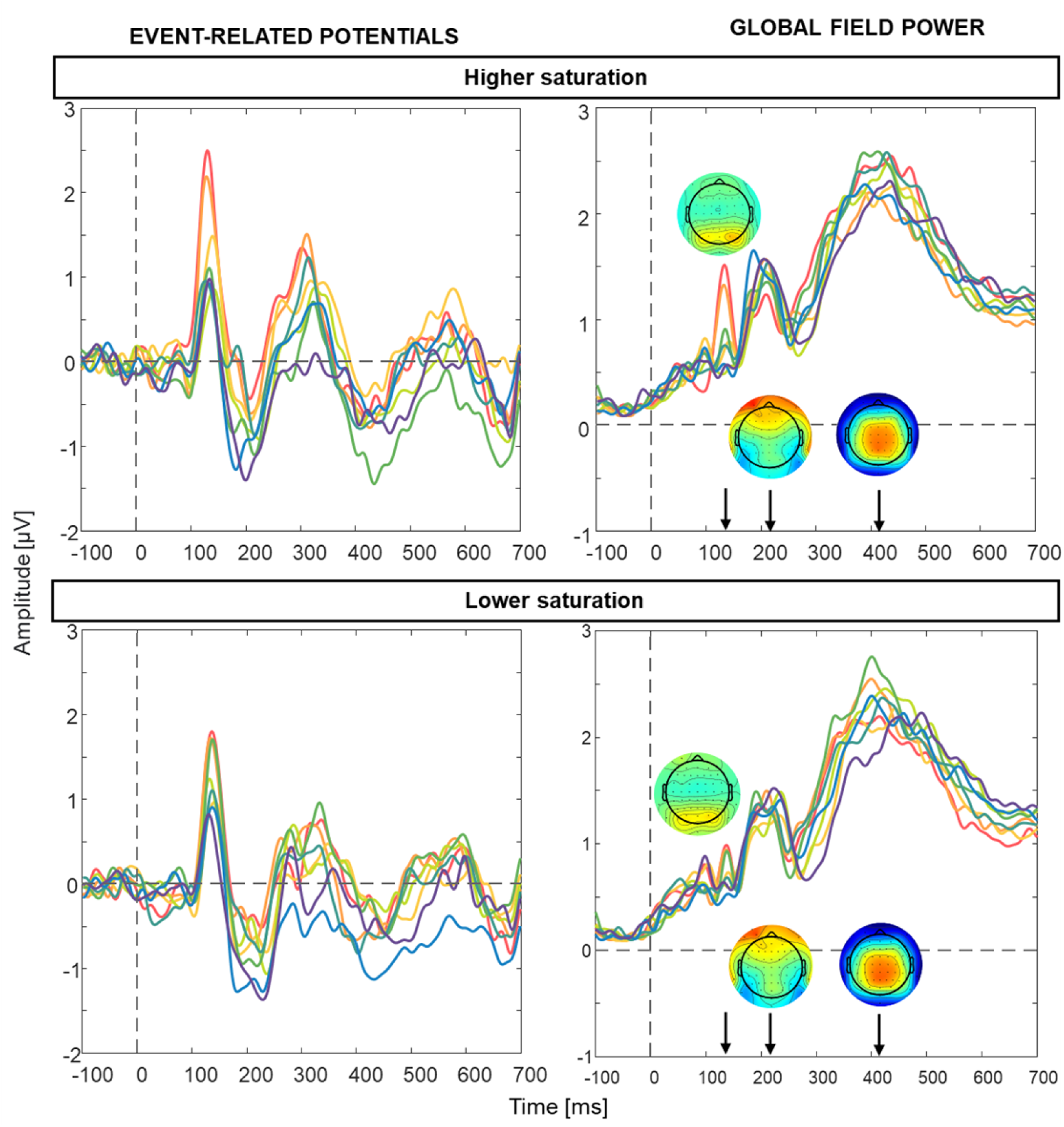
Grand-mean VEPs (left panels) and GFPs (right panel) at occipital sites. Higher saturation is depicted in the top row, with lower saturation below. Note the differences in amplitudes and latencies of the P1 and N1 peaks between hues at different saturation levels – for example, waveforms for orange and red are extremely similar, and waveforms for green appear to be closest to the centre of the distribution, which would make them more similar to their neighbours on either cite in the decoding manifold.

Overall, purple had the highest decoding success in our dataset. This is the case in all sets and across levels of saturation. Neither opponency, nor L or M-cone content in the stimulus provide an explanation for this effect. The S-cone content in the purple stimulus is comparable to that in the turquoise stimulus and second highest in the set, following blue, and is thus an unlikely driver of this effect (see also RSA in Supplementary Materials). The hue scaling content for our purple stimuli is most in line with that of an intermediate hue, with an equal split between red and blue. However, prediction of decoding based on unique hue content would predict better decoding of hues with higher alignment to unique hues, which is not the case for purple, and, as outlined above, seems to be a less likely explanation for decoding within our set compared to the explanations based on low-level content in the stimuli. An alternative high-level explanation could however be a closer proximity to a categorical prototype for purple compared to the other hues. This was however not explicitly measured in our experiment, although higher saturation colours should be closer to prototypes, yet their decoding is generally worse, so this cannot be presumed to be a likely explanation either.

Rosenthal et al. (2021) and Hermann et al. (2022) used stimuli based on intermediate directions between cone-opponent axes and presented them on either a positive or a negative luminance polarity pedestal. Whilst their decoding performance is not directly comparable to the work presented here, an inspection of their model labelling confusion output reveals high frequencies of classification mistakes made between two hue polarities from the same axes, in line with our findings. Rosenthal et al. (2021) and Hermann et al. (2022) also state that decoding is better for stimuli away from the daylight locus axis (i.e., pink and green, compared to blue and yellow). However, higher decoding efficiency may be confounded with the differences in how different cardinal and intermediate chromatic mechanisms integrate with luminance information (Duncan et al., 2012; Martinovic & Andersen, 2018) and the marked influence this could have on their chromatic VEP signature, which would influence decoding performance (Martinovic, 2022). Activations in V1, including those of cone-opponent cells, have been shown to influence chromatic VEP waveforms (Nunez et al., 2021, 2022), with likely impact on representational similarity through amplitude and latency shifts. Our Representational Similarity Analysis (Supplementary Figure 3) shows similar ranges of rank-correlations for both perceptual and lower-level signals – suggesting that decoding results are as likely to be explained by lower-level signals as by perceptual hue-angle differences (for convergent findings, see Kaneko et al., 2020). Putting it all together, it appears that EEG signals do not reflect colour representations that could be easily linked with perceptual or categorical representations derived from behavioural judgments. They are instead largely driven by contrast, so any study aiming to assess high-level representations from EEG activity would need to meticulously control for a myriad of low-level confounds with large impact on waveforms to enable any putative higher-level signatures to become identifiable.

## Acknowledgments

AR was funded by a BBSRS Eastbio PhD studentship (BB/M010996/1) to conduct her research under the supervision of JM.

## Supplementary materials

### Part 1. Supplementary analyses of EEG data

#### 1.1 Pairwise decoding

To verify the findings obtained by decoding hue quadruplets, we also decoded colour pairs. This should provide convergent evidence, both for distant and neighbouring hues. The decoding sets are depicted in Supplementary Figure 1.1., which parallels Figure 4 in the manuscript.

**Supplementary figure 1.1.**
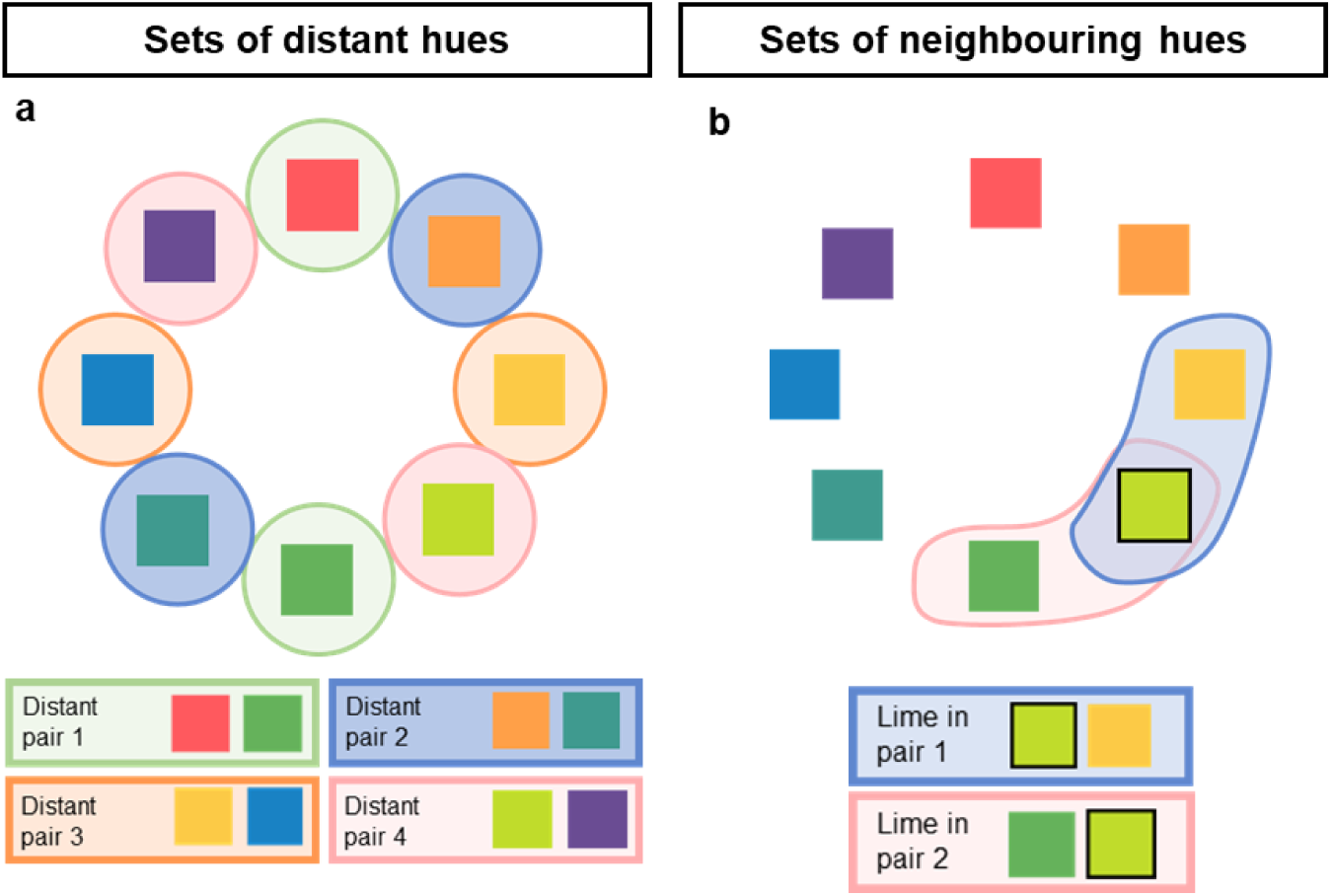
Information Decoding Pairs: Distant and neighbouring hues. On the left (a) the sets examining distant relationships between hues are presented. Distant pairs are decoded in 4 comparisons of opposite hues (e.g., red vs. green). On the right (b) the sets where neighbouring hues are decoded are given. Each hue is presented in two neighbouring pairs, i.e., with each of its neighbours.

##### 1.1.1 EEG decoding: Distant pairs

High mislabelling rates were found between opposite hues in distant quadruplet decoding. This was confirmed when decoding distant hue pairs: accurate decoding was not possible when the signal for each hue was compared only to its opposite hue (Suppl. Figure 1.2).

Note the slight differences in decoding between the two colours in Suppl. Figure 1.2. In tECOC models, the number of LDA units used to classify the data matches the number of labels. This is in contrast to regular binary classifier where, in case of two hues, only a single classifier would be used. In the case of tECOC, probability is combined across separate LDAs. When addressing a binary problem this can lead to asymmetries in probabilities between two labels, especially when the signals related to the two labels are similar, as is evident in Suppl. Figure 1.1. Whilst this indicates tECOC models might not be optimal for resolving binary problems, they remain very efficient and superior to binary classifiers of the One-vs-One and One-vs-All type for problems with more than two classes. For better consistency we use tECOC models for all problems, including pairwise (i.e., binary) hue comparisons.

**Supplementary Figure 1.2.**
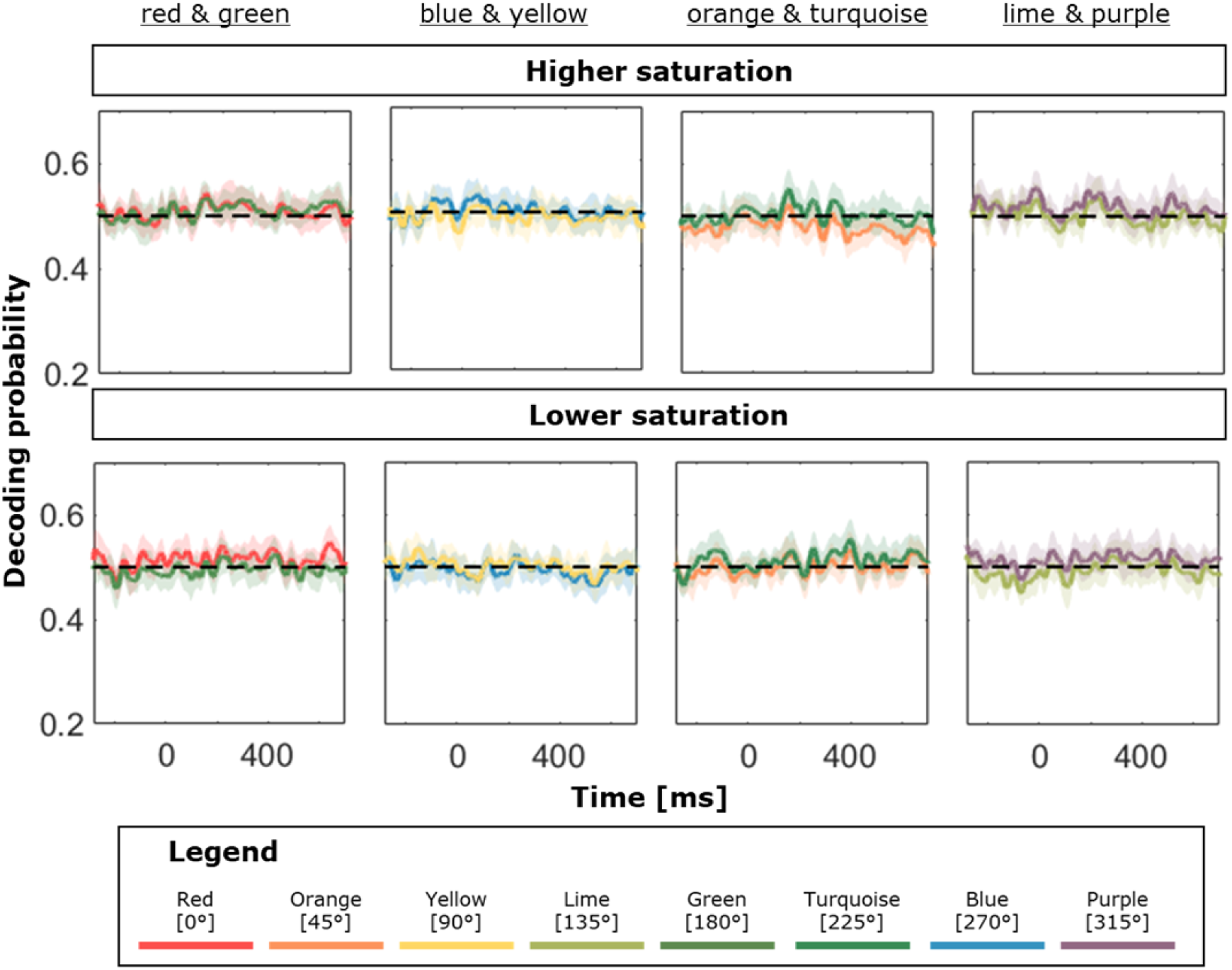
Decoding Accuracy for Distant Pairs. Significant decoding was not possible for any hue in any of the pairs. Error envelopes show 95% confidence intervals.

Taken together, findings from distant quadruplets and pairs alike revealed an unexpected effect of opponency on cortical representation of isoluminant hues as measured using EEG.

##### 1.1.2 EEG decoding: Neighbouring pairs

Decoding of neighbouring pairs allowed us to obtain converging evidence for neighbouring quadruplets by identifying the degree of confusability between immediately neighbouring hues. Yet again, decoding was less successful at higher compared to lower saturation levels. For higher saturation pairs, there were only two hues that could be successfully decoded: yellow from its orange neighbour and purple from both its blue and red neighbours (see Suppl. Figure 1.3).

**Supplementary Figure 1.3.**
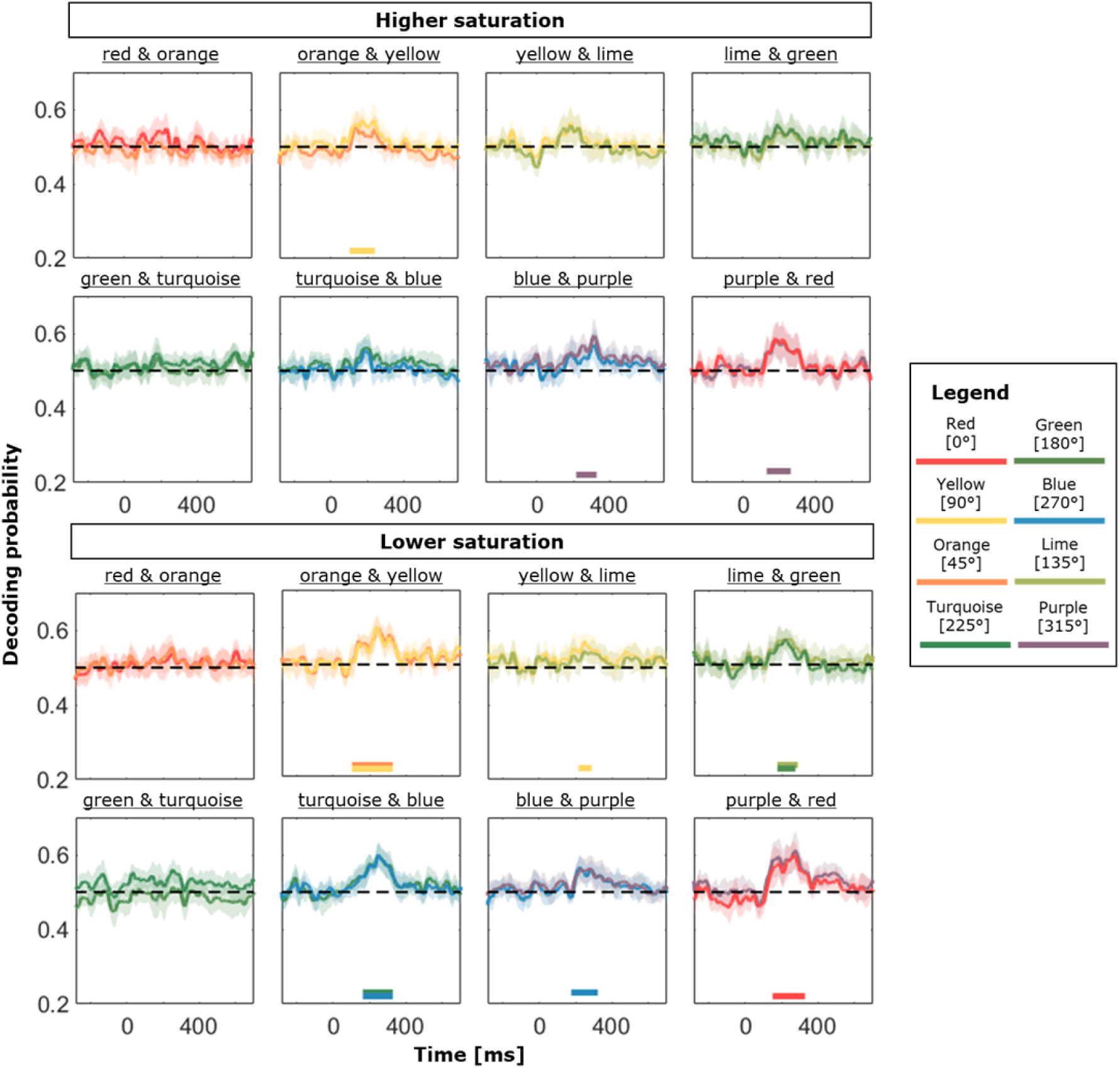
Decoding Accuracy for Neighbouring Pairs. Performance is shown for higher saturation (top panel) and lower saturation (bottom panel). As in other conditions, decoding is better at low saturation levels. Red and orange, and green and turquoise cannot be decoded from each other at either saturation level. Error envelopes show 95% confidence intervals. The lines at the bottom indicate significant decoding from chance level (50%) in the 100-300 ms window.

At lower saturation levels, orange and yellow were significantly decoded from each other, and so were lime and green and turquoise and blue. Blue was also decoded with significant success from its purple neighbour. This was the case for purple and red too.

Fitting a linear mixed effect model to decoding accuracy between 100 and 300 ms (see Suppl. Figure 1.4), we found no interaction (χ2(7)=7.730, p = .357), but rather an additive effect of saturation (χ2(1)=7.077, p-p=.008) and colour (χ2(7)=22.566, p=.002). This is different to the analysis of neighbouring quadruplets, where the two factors interacted - it is plausible that the lack of interaction is driven by the fact that only one representational overlap is affecting decoding performance, rather than the overlap with representations of three other colours (as in quadruplets). The analysis on quadruplets also has twice as many datapoints and is capable of explaining approx. twice as much variance, both for fixed and random effects. Despite those concerns, the overall pattern of results is largely similar: green is decoded somewhat worse and purple somewhat better, with better decoding at lower saturation.

**Supplementary Figure 1.4.**
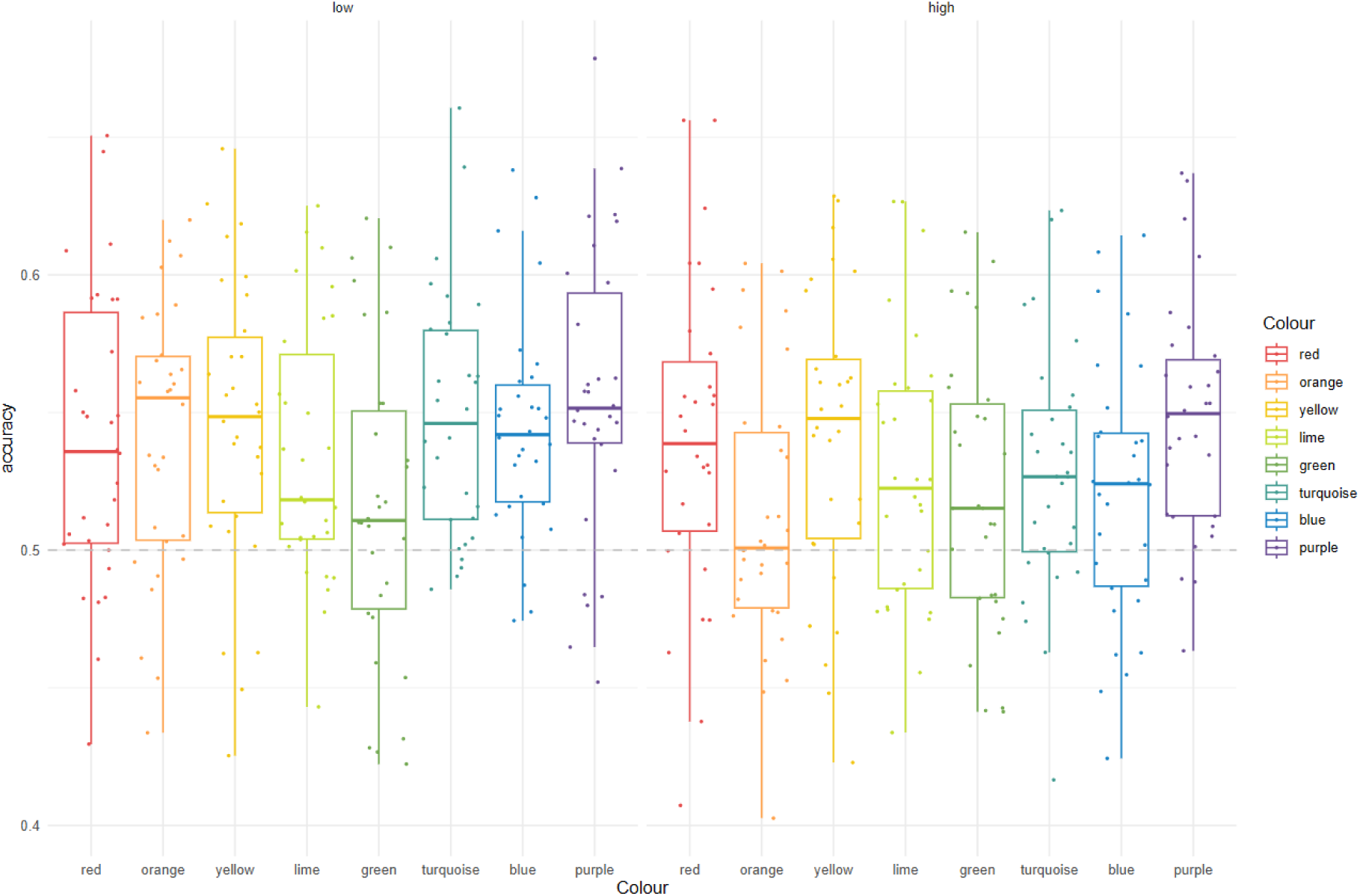
Box plot of accurate decoding proportions over the 100-300 ms window for neighbouring pairs. *Horizontal lines are medians,* b*oxes encompass 50% of the distribution (the interquartile range – IQR), the minimum and maximum whisker values extend the box by -/+ 1.5 * IQR to depict variability in each direction, while dots show individual participants*.

Horizontal dashed line at 0.5 denotes the chance decoding level. Note that decoding is (1) highest for purple, (2) lowest for green, and (3) lower for high saturation colours. It is useful to compare this figure with Figure 14 from the manuscript, depicting accurate decoding of neighbouring quadruplets.

#### 1.2 EEG decoding: Convergence of pairwise decoding with Difference Wave Analyses

Pairwise Decoding using LDA is equivalent to training the classifier to place a boundary midway through subsampled sets of difference waves (with each set corresponding to one decoding fold) in a multidimensional electrode space. In other words, the position of the decision boundary used for classifying the training sets linearly depends on the size of amplitude differences in voltage across the electrode set. This means that information decoding with LDA is simply a much powerful way of doing standard ERP analyses - a multivariate approach to inherently multivariate (i.e., multi-electrode) data, instead of a univariate approach imposed by averaging across electrodes in a cluster as used in standard ERPs.

When decoding colour from spatially identical patterns (here, circles) it is assumed that the dipoles driving EEG activity will largely reflect the differential activation of colour-processing networks. Chauhan et al. (2023) demonstrated that 8 occipital channels are sufficient for classification performance in study using similar stimulus properties. In a sense, this is a validation of the standard visual evoked potential technique, which relies on the assumption that averaged activity over occipital electrodes is a valid representation of the stimulus-elicited visuocortical activity.

In this supplementary section, we demonstrate that for our dataset, a substantial portion of the decoding variance can be explained by ERP fluctuations at the occipital electrode cluster (Suppl. Figure 1.5 – decoding yellow and orange). First, we calculate a grand-mean difference wave by subtracting amplitude for yellow from amplitude for orange. Then, we take the absolute value of differential amplitude, as only the magnitude of the difference rather than its direction is relevant for the decoder. We then correlate the absolute value of the difference wave with decoding accuracy across the samples encompassing the period between -100 and 600 ms (r(718) = 0.486, p < .001). While a non-negligible proportion of the variance across time (r2=0.236) can be explained by difference wave properties, some important information is lost – this should not come as a surprise, since differences between occipital electrodes are removed through averaging, while other electrodes are left out of the occipital cluster altogether.

**Supplementary Figure 1.5.**
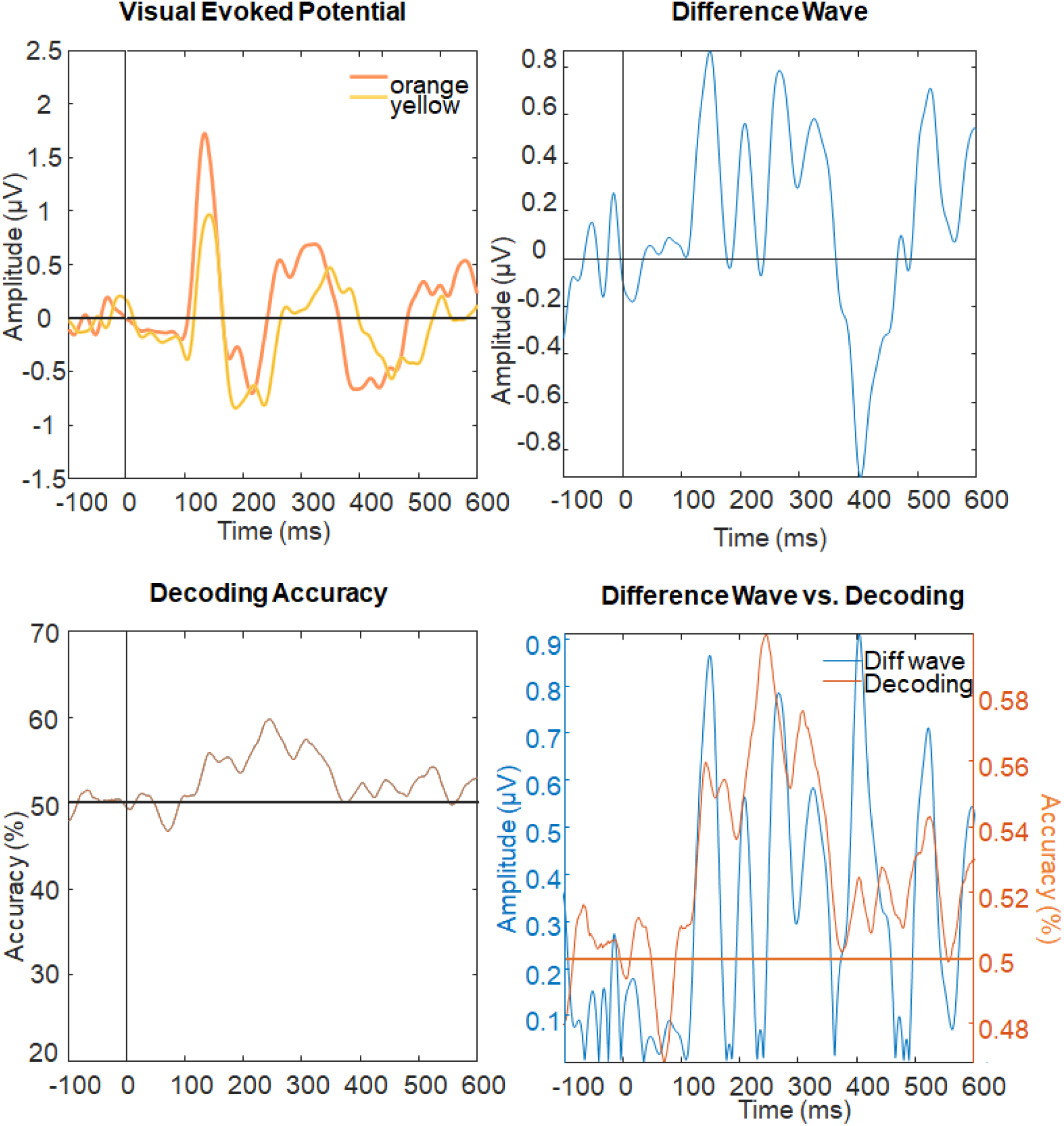
Liking decoding accuracy to difference waves. Top left panel: Grand-mean visual evoked potentials for orange and yellow at the occipital electrode cluster. Top right panel: Difference wave, obtained by subtracting yellow from orange amplitude depicted in the top left panel. Bottom left panel: Decoding accuracy for yellow and orange, averaged across the two LDA units. Bottom right panel: in blue, the absolute value of the difference wave is depicted; in orange, decoding accuracy is depicted. Note that the decoding accuracy matches the fluctuations in the difference wave, with the rise in accuracy matching the increase in differences observed in ∼100-300 ms.

### Part 2. Statistical Models

2.1 Best fitting generalised linear mixed effect model for accuracy binomial data from oddball targets, with random intercepts only

| **Fixed Effects** | **Estimate ±**<br>**SE** | **z**<br>**value** | **p value** | **Odds**<br>**ratio,**<br>**(95% CI)** |
| --- | --- | --- | --- | --- |
| Intercept | 1.955 ±<br>0.120 | 16.32 | <.001*** | 7.06<br>(5.59,<br>8.93) |

Random effects: The maximal model that could be adequately fitted to the dataset included only random by-participant intercepts, which accounted for 0.155 ± 0.393 in variance (variance ± SD). Residual variance 3.29 (ICC = 0.04). There were 2400 responses analysed, from a total of 15 participants. Conditional r2: 0.045; Marginal r2: 0.

2.2 Best fitting linear mixed effect model for reaction time on hits, with dummy coding for hue (red, orange, yellow, lime, green, turquoise, blue and purple) and saturation (low and high).

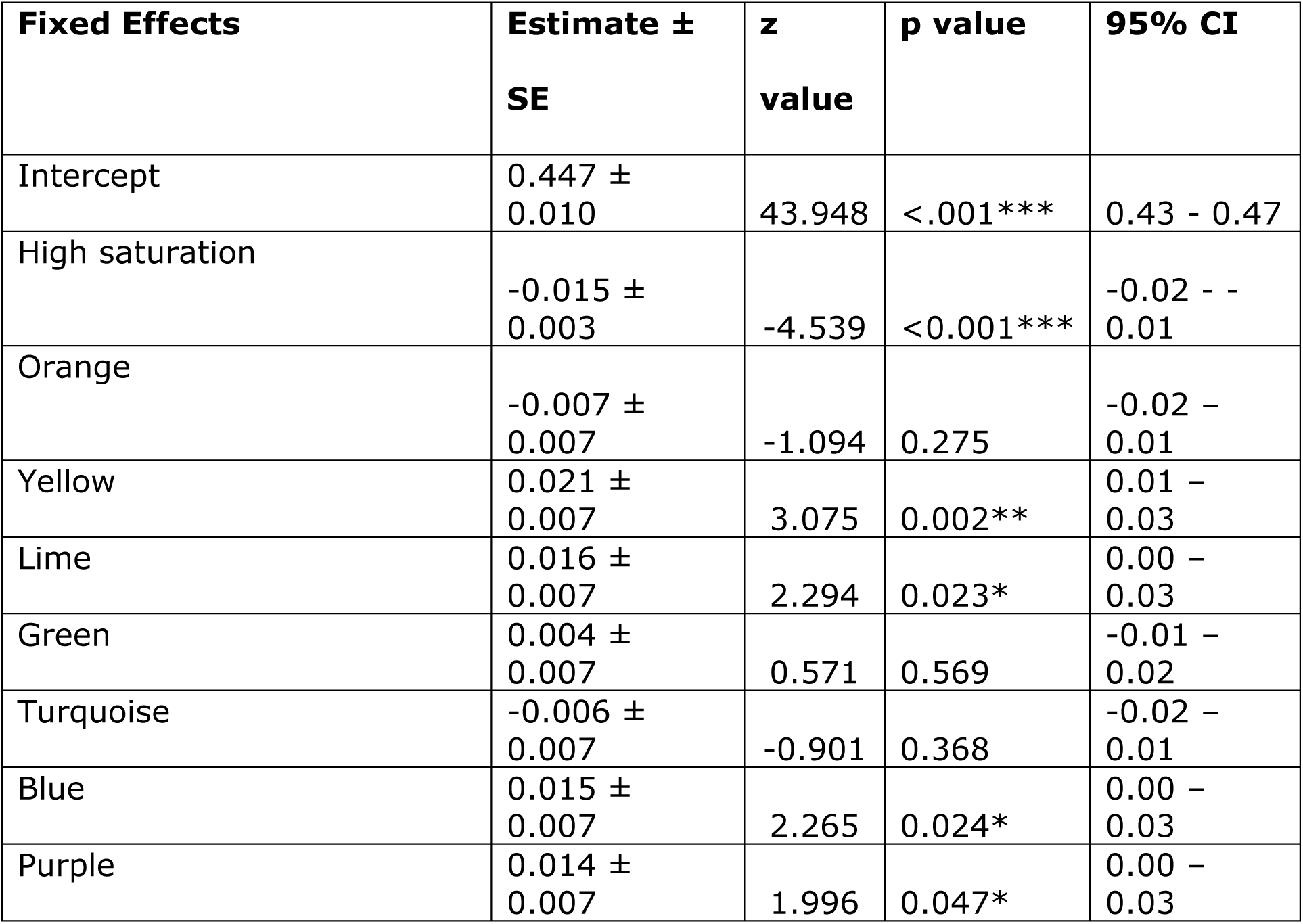
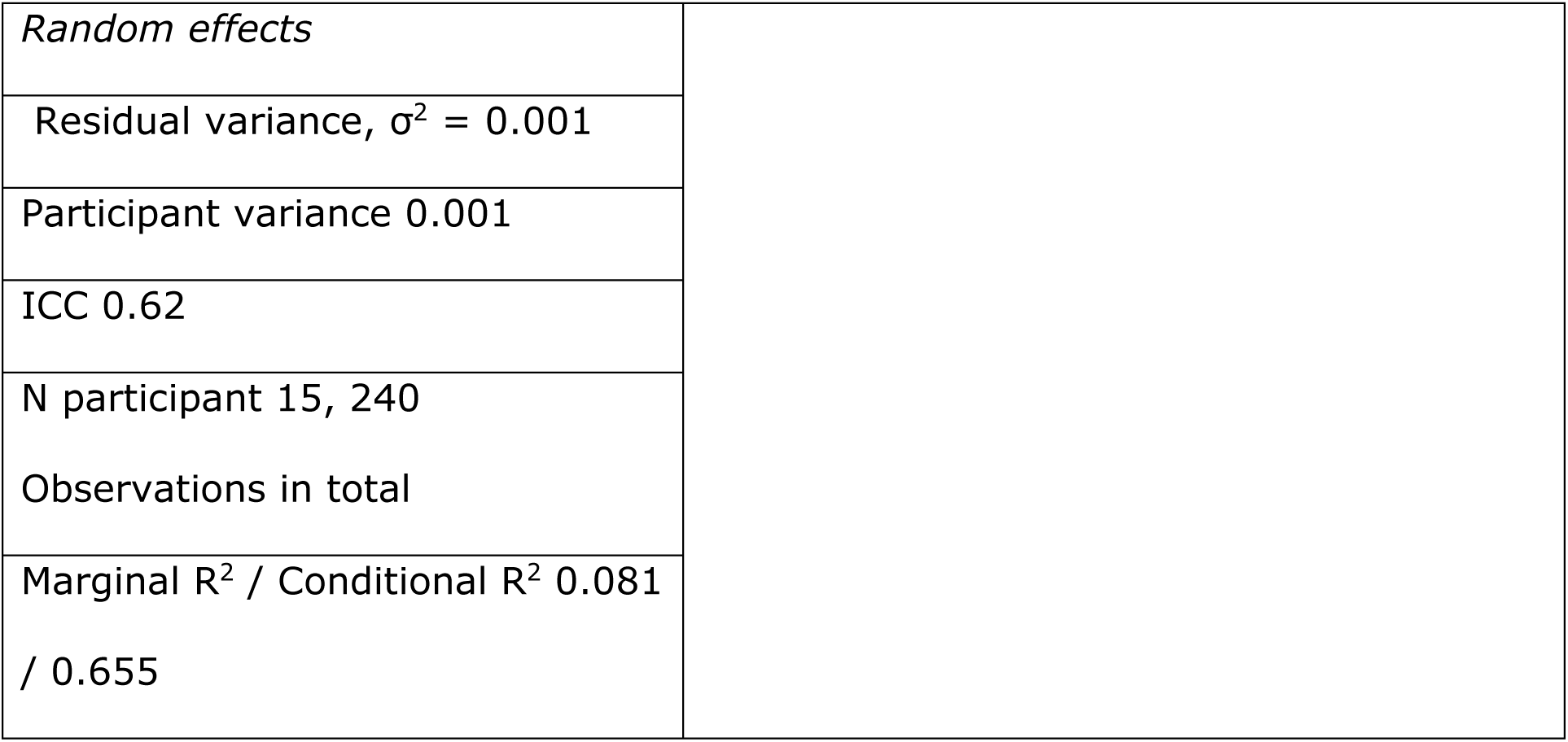

| **Fixed Effects** | **Estimate ±**<br>**SE** | **z**<br>**value** | **p value** | **95% CI** |
| --- | --- | --- | --- | --- |
| Intercept | 0.447 ±<br>0.010 | 43.948 | <.001*** | 0.43 - 0.47 |
| High saturation | -0.015 ±<br>0.003 | -4.539 | <0.001*** | -0.02 - -<br>0.01 |
| Orange | -0.007 ±<br>0.007 | -1.094 | 0.275 | -0.02 -<br>0.01 |
| Yellow | 0.021 ±<br>0.007 | 3.075 | 0.002** | 0.01 -<br>0.03 |
| Lime | 0.016 ±<br>0.007 | 2.294 | 0.023* | 0.00 -<br>0.03 |
| Green | 0.004 ±<br>0.007 | 0.571 | 0.569 | -0.01 -<br>0.02 |
| Turquoise | -0.006 ±<br>0.007 | -0.901 | 0.368 | -0.02 -<br>0.01 |
| Blue | 0.015 ±<br>0.007 | 2.265 | 0.024* | 0.00 -<br>0.03 |
| Purple | 0.014 ±<br>0.007 | 1.996 | 0.047* | 0.00 -<br>0.03 |

|  |  |
| --- | --- |
| *Random effects* |  |
| Residual variance, $\sigma^2 = 0.001$ | |
| Participant variance 0.001 |  |
| ICC 0.62 |  |
| N participant 15, 240<br>Observations in total |  |
| Marginal $R^2$ / Conditional $R^2$ 0.081<br>/ 0.655 | |

2.3 Best fitting linear mixed effect model for hue scaling of redness, with fixed effect of hue (0° - baseline, 45° and 315°; (2)= 41.195, p<.001) and random intercepts only.

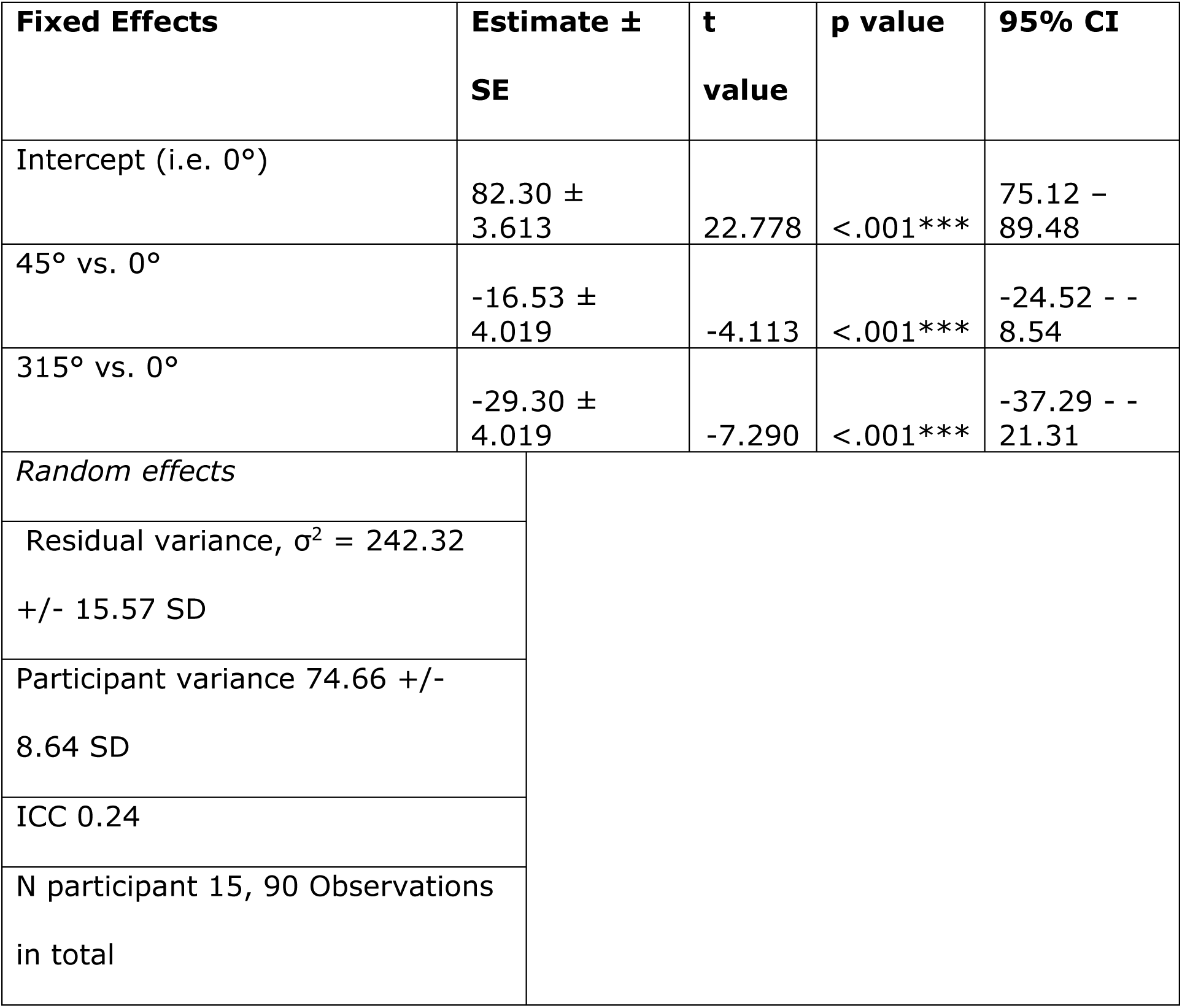
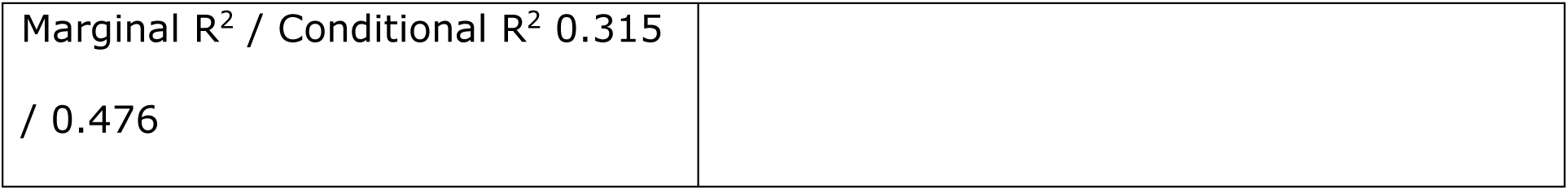

| **Fixed Effects** | **Estimate ±**<br>**SE** | **t**<br>**value** | **p value** | **95% CI** |
| --- | --- | --- | --- | --- |
| Intercept (i.e. 0°) | 82.30 ±<br>3.613 | 22.778 | <.001*** | 75.12 -<br>89.48 |
| 45° vs. 0° | -16.53 ±<br>4.019 | -4.113 | <.001*** | -24.52 - -<br>8.54 |
| 315° vs. 0° | -29.30 ±<br>4.019 | -7.290 | <.001*** | -37.29 - -<br>21.31 |
| *Random effects* |  |  |  |  |
| Residual variance, $\sigma^2 = 242.32$<br>+/- 15.57 SD | | | | |
| Participant variance 74.66 +/-<br>8.64 SD |  |  |  |  |
| ICC 0.24 |  |  |  |  |
| N participant 15, 90 Observations<br>in total |  |  |  |  |

|  |
| --- |
| Marginal R<sup>2</sup> / Conditional R<sup>2</sup> 0.315<br>/ 0.476 |

Fixed effects that did not contribute to the model: interaction χ2(2)=3.497, p=.174; chroma χ2(1)=0.273, p=.602. Post-hoc paired t-test indicates that besides the significant differences for both orange and purple against red, there is also a significant difference between orange and purple of 12.8% +/- 4.02% SE, t(73)=3.176, p=.0062.

2.4 Best fitting linear mixed effect model for hue scaling of greenness, with fixed effect of hue (180° - baseline, 135° and 225°; =118.06, p<.001) and random intercepts only.

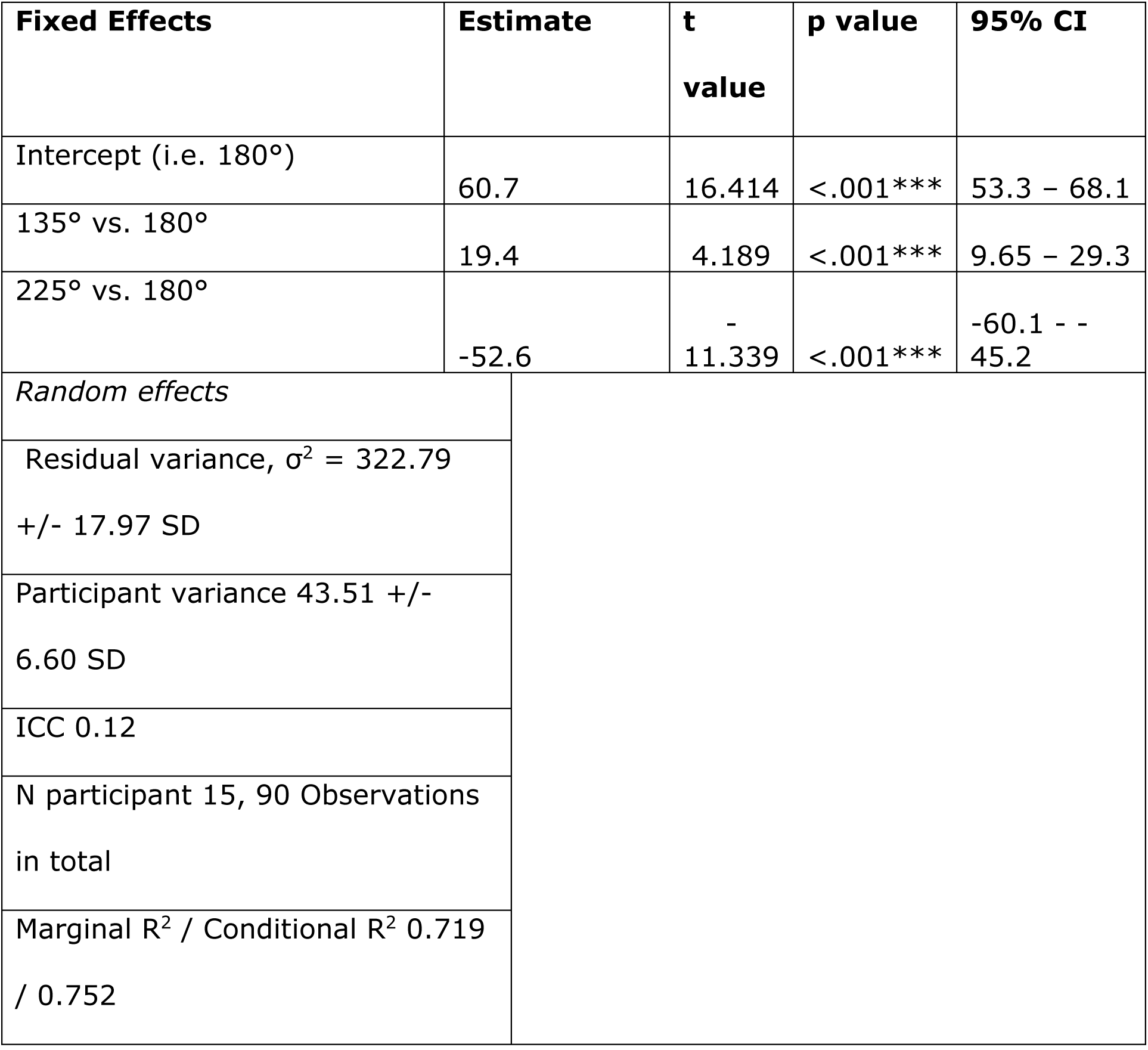

| Fixed Effects | Estimate | t value | p value | 95% CI |
| --- | --- | --- | --- | --- |
| Intercept (i.e. 180°) | 60.7 | 16.414 | <.001*** | 53.3 – 68.1 |
| 135° vs. 180° | 19.4 | 4.189 | <.001*** | 9.65 – 29.3 |
| 225° vs. 180° | -52.6 | -11.339 | <.001*** | -60.1 - -45.2 |
| *Random effects* |  |  |  |  |
| Residual variance, $\sigma^2 = 322.79$<br>+/- 17.97 SD | | | | |
| Participant variance 43.51 +/- 6.60 SD |  |  |  |  |
| ICC 0.12 |  |  |  |  |
| N participant 15, 90 Observations in total |  |  |  |  |
| Marginal R<sup>2</sup> / Conditional R<sup>2</sup> 0.719<br>/ 0.752 |  |  |  |  |

Fixed effects that did not contribute to the model: interaction χ2(2)=3.999, p=.135; chroma χ2(1)=0.811, p=.368. Post-hoc paired t-test indicates that besides the significant differences for both lime and turquoise against green, there is also a significant difference between lime and turquoise of 72.0% +/- 4.64% SE, t(73)=15.528, p<.001. Note: evaluation of model fit revealed non-homogenous variance, which is why we report estimates and 95% CIs from a bootstrapped model (1000 resamples of participants). We report the statistical tests for the parametric model.

2.5 Best fitting linear mixed effect model for hue scaling of blueness, with fixed effect of hue (270° - baseline, 180°, 225° and 315°) and random intercepts only.

| **Fixed Effects** | **Estimate ±**<br>**SE** | **t**<br>**value** | **p value** | **95% CI** |
| --- | --- | --- | --- | --- |
| Intercept (i.e. 270°) | 89.6 | 24.448 | <.001*** | 82.1 – 96.9 |
| 180° vs. 270° | -56.2 | -<br>12.148 | <.001*** | -68.6 - -<br>43.4 |
| 225° vs. 270° | -1.80 | -0.389 | 0.698 | -10.6 -<br>7.28 |
| 315° vs. 270° | -43.4 | -9.375 | <.001*** | -53.9 - -<br>32.5 |
| *Random effects* |  |  |  |  |
| Residual variance, $\sigma^2 = 321.43$<br>+/- 17.93 SD | | | | |
| Participant variance 40.60 +/-<br>6.37 SD |  |  |  |  |
| ICC 0.11 |  |  |  |  |
| N participant 15, 120 Observations<br>in total |  |  |  |  |
| Marginal $R^2$ / Conditional $R^2$ 0.633<br>/ 0.674 | | | | |

Fixed effects that did not contribute to the model: interaction χ2(3)=1.9032, p=.593; chroma χ2(1)=0.006, p=.938. Post-hoc paired t-test indicates that besides the significant differences for both green and purple against blue, there is also a significant difference between purple and turquoise of 41.6% +/- 4.63% SE, t(102)=8.987, p<.001. Note: evaluation of model fit revealed non-homogenous variance and non-normally distributed residuals, which is why we report estimates and 95% CIs from a bootstrapped model (1000 resamples of participants). We report the statistical tests for the parametric model.

2.6 Best fitting linear mixed effect model for hue scaling of yellowness, with fixed effect of hue (90° - baseline, 45° and 135°, (2)= 114.82, p<.001) and random intercepts only.

| **Fixed Effects** | **Estimate ±**<br>**SE** | **t**<br>**value** | **p value** | **95% CI** |
| --- | --- | --- | --- | --- |
| Intercept (i.e. 90°) | 86.23 ±<br>3.347 | 25.76 | <.001*** | 79.58 –<br>92.89 |
| 45° vs. 90° | -53.67 ±<br>4.700 | -11.42 | <.001*** | -63.01 – -<br>44.32 |
| 135° vs. 90° | -66.67 ±<br>4.700 | -14.19 | <.001*** | -76.01 – -<br>57.32 |
| *Random effects* |  |  |  |  |
| Residual variance, $\sigma^2 = 331.32$<br>+/- 18.20 SD | | | | |
| Participant variance 2.42 +/- 1.56<br>SD |  |  |  |  |
| ICC 0.01 |  |  |  |  |
| N participant 15, 90 Observations<br>in total |  |  |  |  |
| Marginal $R^2$ / Conditional $R^2$ 0.716<br>/ 0.718 | | | | |

Fixed effects that did not contribute to the model: interaction χ2(2)=3.253, p=.197; chroma χ2(1)=1.640, p=.200. Post-hoc paired t-test indicates that besides the significant differences for both orange and lime against yellow, there is also a significant difference between orange and lime themselves, with orange being perceived as yellower 13.0% +/- 4.7% SE, t(73)=2.766, p=.0194.

2.7 Best fitting linear mixed effect model for decoding accuracy for distant quadruplets, with fixed effects of hue (0° - baseline, 45°, 90°, 135°, 180°, 225°, 270° and 315°) and saturation (low and high) and random intercepts only.

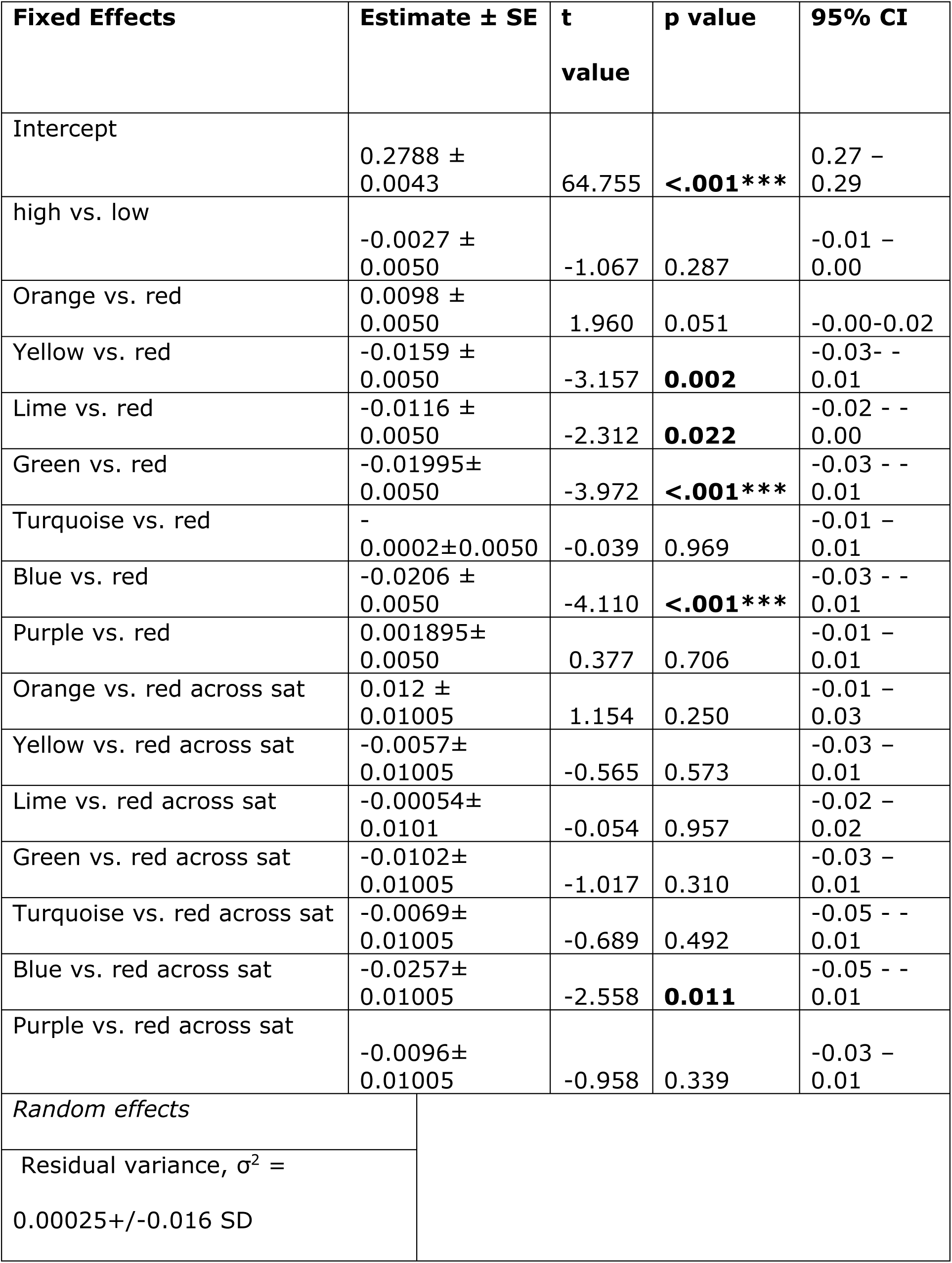
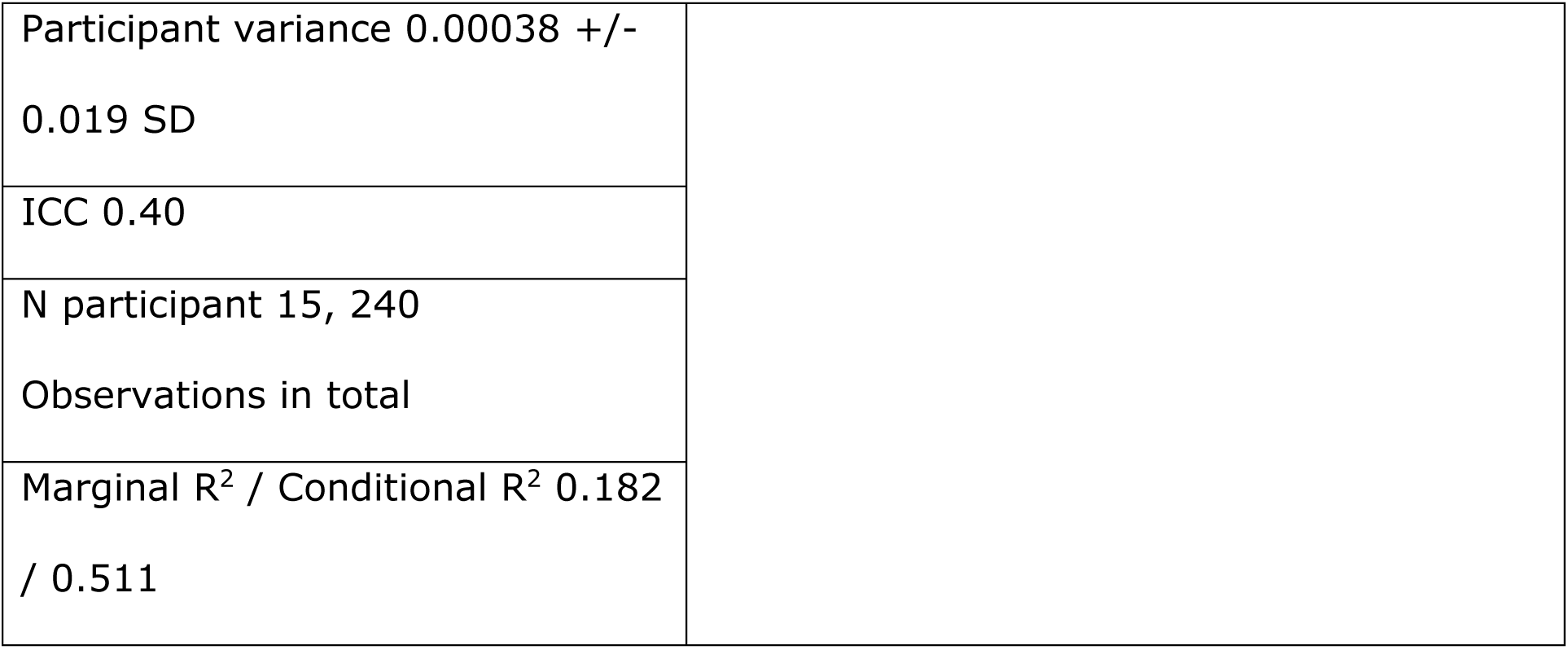

| Fixed Effects | Estimate ± SE | t<br>value | p value | 95% CI |
| --- | --- | --- | --- | --- |
| Intercept | 0.2788 ± 0.0043 | 64.755 | **<.001\*\*\*** | 0.27 – 0.29 |
| high vs. low | -0.0027 ± 0.0050 | -1.067 | 0.287 | -0.01 – 0.00 |
| Orange vs. red | 0.0098 ± 0.0050 | 1.960 | 0.051 | -0.00-0.02 |
| Yellow vs. red | -0.0159 ± 0.0050 | -3.157 | **0.002** | -0.03- - 0.01 |
| Lime vs. red | -0.0116 ± 0.0050 | -2.312 | **0.022** | -0.02 - - 0.00 |
| Green vs. red | -0.01995± 0.0050 | -3.972 | **<.001\*\*\*** | -0.03 - - 0.01 |
| Turquoise vs. red | - 0.0002±0.0050 | -0.039 | 0.969 | -0.01 – 0.01 |
| Blue vs. red | -0.0206 ± 0.0050 | -4.110 | **<.001\*\*\*** | -0.03 - - 0.01 |
| Purple vs. red | 0.001895± 0.0050 | 0.377 | 0.706 | -0.01 – 0.01 |
| Orange vs. red across sat | 0.012 ± 0.01005 | 1.154 | 0.250 | -0.01 – 0.03 |
| Yellow vs. red across sat | -0.0057± 0.01005 | -0.565 | 0.573 | -0.03 – 0.01 |
| Lime vs. red across sat | -0.00054± 0.0101 | -0.054 | 0.957 | -0.02 – 0.02 |
| Green vs. red across sat | -0.0102± 0.01005 | -1.017 | 0.310 | -0.03 – 0.01 |
| Turquoise vs. red across sat | -0.0069± 0.01005 | -0.689 | 0.492 | -0.05 - - 0.01 |
| Blue vs. red across sat | -0.0257± 0.01005 | -2.558 | **0.011** | -0.05 - - 0.01 |
| Purple vs. red across sat | -0.0096± 0.01005 | -0.958 | 0.339 | -0.03 – 0.01 |
| Random effects |  |  |  |  |
| Residual variance, σ² = 0.00025+/-0.016 SD |  |  |  |  |

|  |
| --- |
| Participant variance 0.00038 +/-<br>0.019 SD |
| ICC 0.40 |
| N participant 15, 240<br>Observations in total |
| Marginal R<sup>2</sup> / Conditional R<sup>2</sup> 0.182<br>/ 0.511 |

There was a significant interaction between saturation and hue, χ2(7)=15.232, p=.033. Post-hoc paired t-tests indicate that high saturation orange is decoded better than high (t t(241)=4.670, p<.001) and low (t(241)=4.334, p=.002) saturation yellow, high (t(241)=3.744, p=0.02) and low (t(241) = 4.106, p=.005) saturation lime, high (t(241)=5.536, p<.001) and low (t(241) = 4.582, p<.001) saturation green, and high (t(241)=6.682, p<.001) and low (t(241) = 3.623, p =.0301) saturation blue. Meanwhile, the worst decoded colour is high saturation blue – besides being more poorly decoded than high (see above) and low (t(241)=4.670, p<.001) saturation orange, it is also more poorly decoded against high (t(241) = 4.159, p = .005) and low (t(241)=5.031, p<.001) saturation purple, high (t(241) = 4.058, p=.007) and low (t(241) = 4.563, p<.001) saturation turquoise, and high (t(241) = 4.555, p<.001) and low (t(241)=4.119, p=.005) saturation red.

2.8 Best fitting linear mixed effect model for decoding accuracy of neighbouring quadruplets, with fixed effects of hue (0° - baseline, 45°, 90°, 135°, 180°, 225°, 270° and 315°) and saturation (low and high) and random intercepts only.

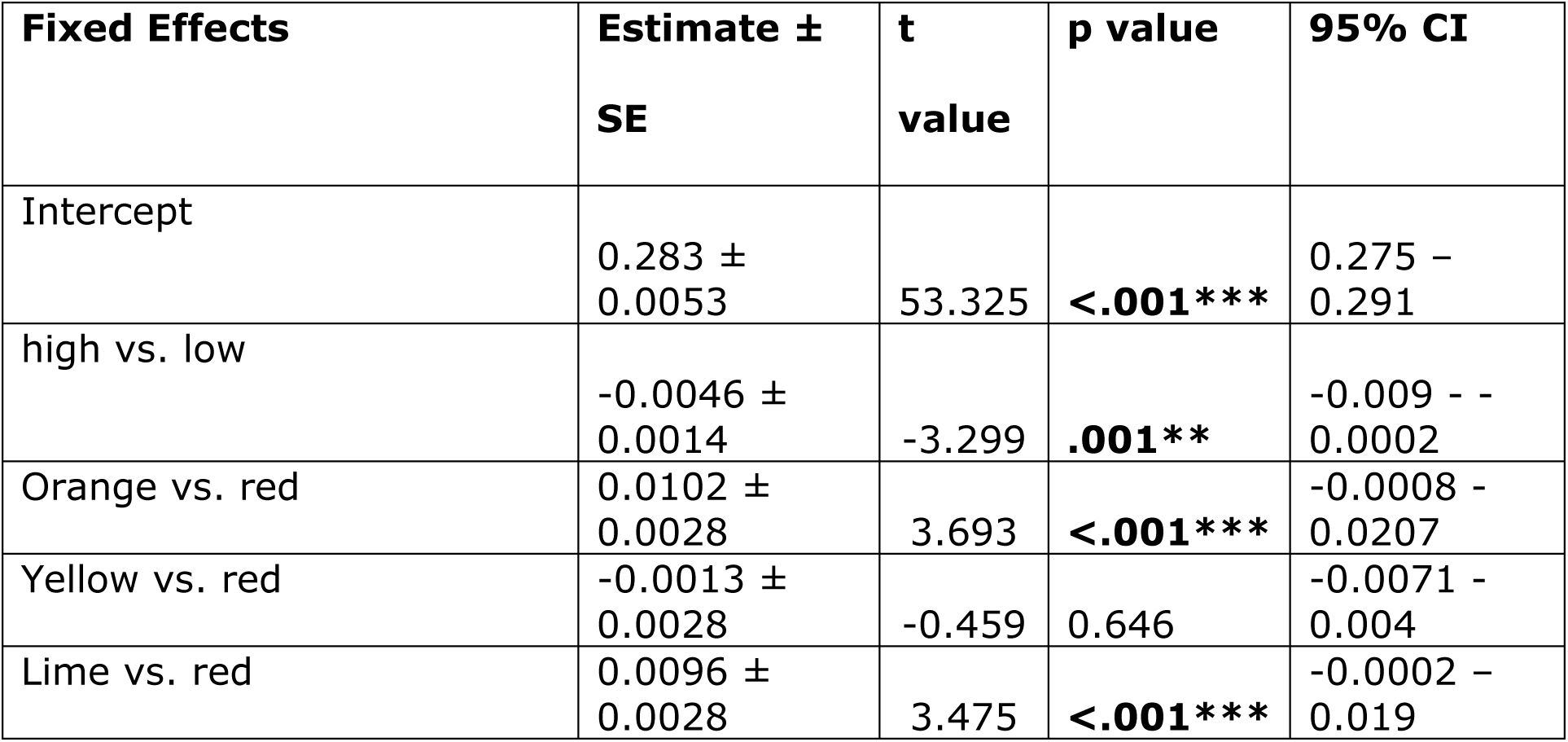
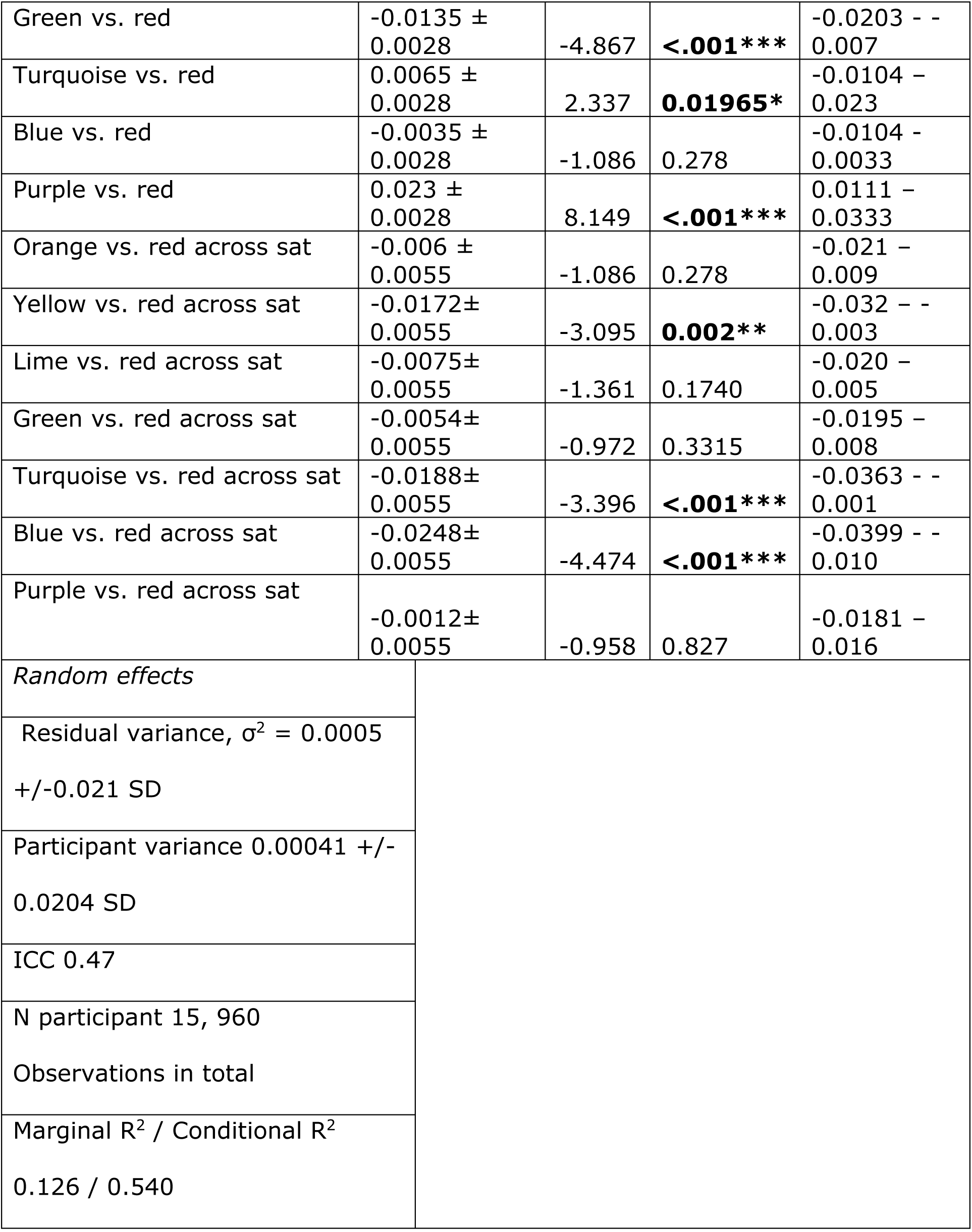

There was a significant interaction between saturation and hue, χ2(7)=36.3, p<.001. Post-hoc paired t-test indicates that decoding is best for purple at higher saturation (ps < .03 for all other colours apart from low saturation turquoise t(960) = 2.946, p = .196 and low saturation purple t(960) = 1.098, p = .999) and poorest for low saturation green (ps <.005 for all other colours apart from high saturation yellow t(960) = 1.643, p = 0.960, high saturation blue t(960) = 0.118, p = 1.00, high saturation turquoise (t(960) = 3.393, p = .058, low saturation red t(960) = 2.691, p=.338 and high saturation green t(960) = 0.041, p=1.00). Colour with a significant drop in decoding across the two levels of saturation is blue (t(960) = 4.873, p<.001).

2.9 Best fitting linear mixed effect model for decoding accuracy of neighbouring pairs, with fixed effects of hue (0° - baseline, 45°, 90°, 135°, 180°, 225°, 270° and 315°) and saturation (low and high) and random intercepts only.

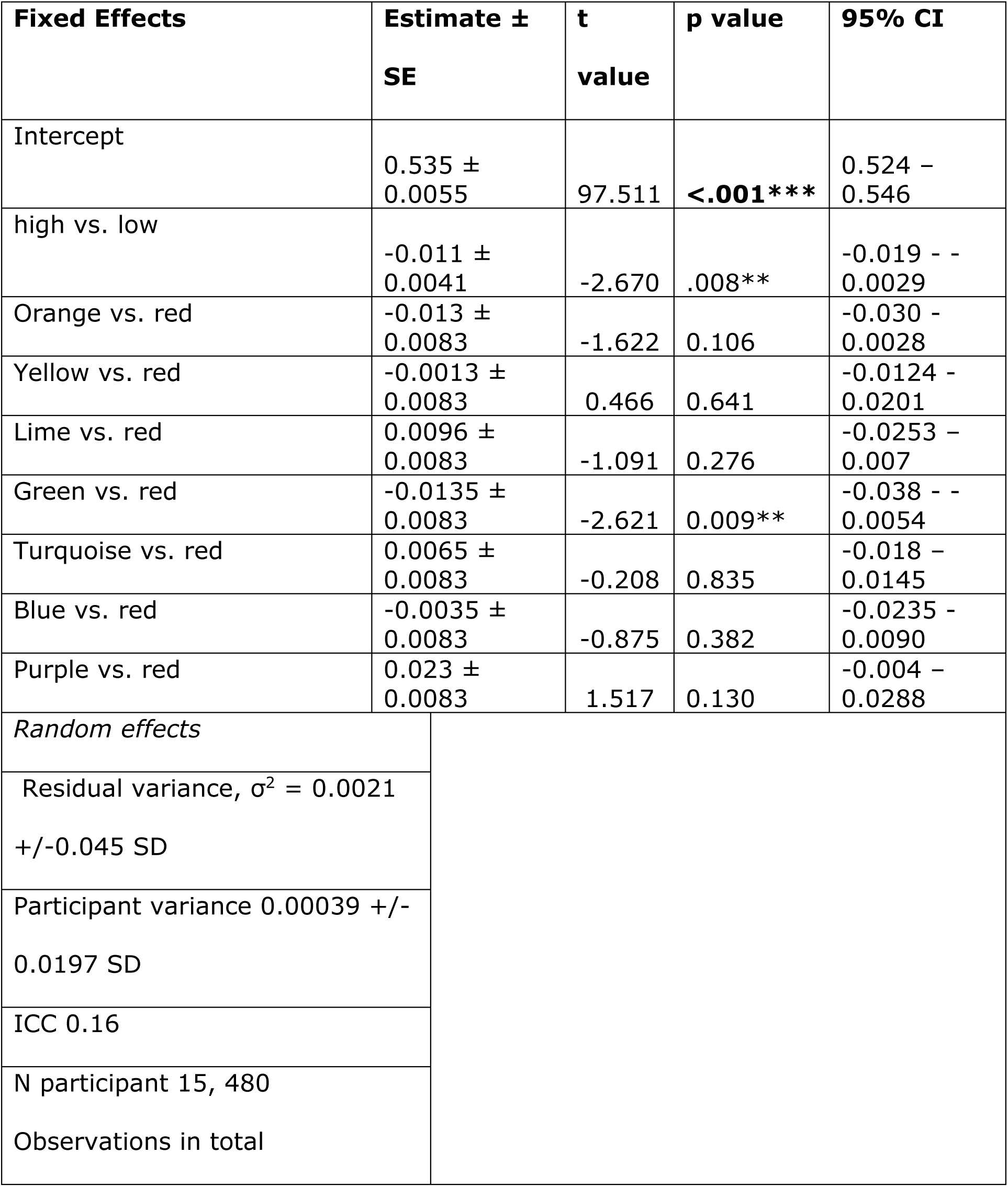
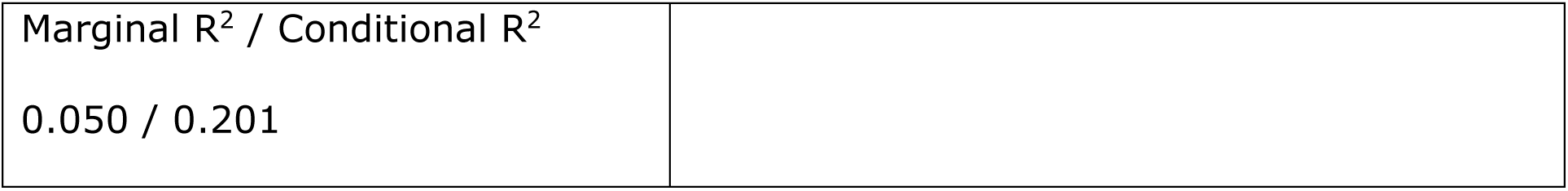

| Fixed Effects | Estimate ±<br>SE | t<br>value | p value | 95% CI |
| --- | --- | --- | --- | --- |
| Intercept | 0.535 ±<br>0.0055 | 97.511 | <.001*** | 0.524 -<br>0.546 |
| high vs. low | -0.011 ±<br>0.0041 | -2.670 | .008** | -0.019 - -<br>0.0029 |
| Orange vs. red | -0.013 ±<br>0.0083 | -1.622 | 0.106 | -0.030 -<br>0.0028 |
| Yellow vs. red | -0.0013 ±<br>0.0083 | 0.466 | 0.641 | -0.0124 -<br>0.0201 |
| Lime vs. red | 0.0096 ±<br>0.0083 | -1.091 | 0.276 | -0.0253 -<br>0.007 |
| Green vs. red | -0.0135 ±<br>0.0083 | -2.621 | 0.009** | -0.038 - -<br>0.0054 |
| Turquoise vs. red | 0.0065 ±<br>0.0083 | -0.208 | 0.835 | -0.018 -<br>0.0145 |
| Blue vs. red | -0.0035 ±<br>0.0083 | -0.875 | 0.382 | -0.0235 -<br>0.0090 |
| Purple vs. red | 0.023 ±<br>0.0083 | 1.517 | 0.130 | -0.004 -<br>0.0288 |
| *Random effects* |  |  |  |  |
| Residual variance, $\sigma^2 = 0.0021$<br>+/-0.045 SD | | | | |
| Participant variance 0.00039 +/-<br>0.0197 SD |  |  |  |  |
| ICC 0.16 |  |  |  |  |
| N participant 15, 480<br>Observations in total |  |  |  |  |

|  |  |
| --- | --- |
| Marginal $R^2$ / Conditional $R^2$ | |
| 0.050 / 0.201 |  |

Post-hoc paired t-test indicate that decoding is better for purple compared to orange t(473) = 3.112, p = .0410 and green t(473) = 4.103, p = .0012) and for yellow compared to green t(473) = 3.061, p=.0476.

### Part 3. Representational Similarity Analysis

For each observer, the decoding scores (a time-series of 4x4 confusion matrices) for each model were used to construct a time-series of representational dissimilarity matrices (RDMs) using the method outlined in Chauhan et al. (2023).

The stimuli for each observer were then used to construct static RDMs using a variety of perceptual and physiological coordinates (see Suppl. Figure 3.1). We used the hue-angle from the LCh coordinates of the stimuli, hue-scaling measurements, LMS cone excitations, cone-contrasts calculated against the grey background, and the opponency signals in DKL coordinates.

**Supplementary Figure 3.1.**
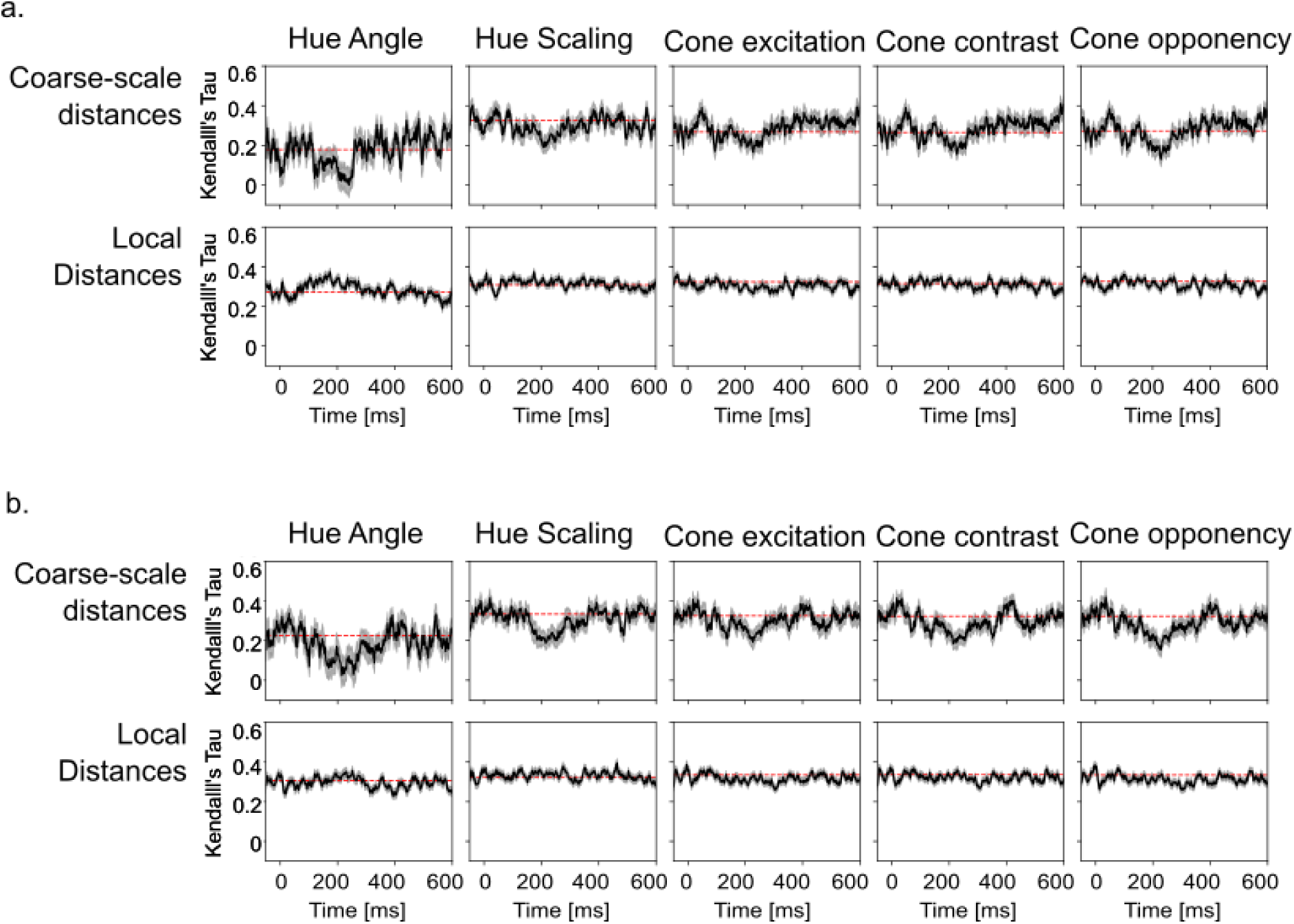
Representational similarity analysis, depicting rank correlation with different perceptual or physiological coordinates. Panel (a) depicts them for high-saturation stimuli, while panel b shows the same for low-saturation stimuli. In all panels, a dashed red line marks the average correlation over 50ms immediately preceding the stimulus.

For each observer, the classifier RDM at each time-point was correlated with the five static RDMs – giving us a time-series of rank-correlations. For both distal and neighbouring-hues cases, we averaged over all models (2 for distal-hues and 8 for neighbouring-hues), and then averaged across all observers. Note that distal-hues represent coarse-scale distances, while neighbouring-hues represent ordinally smaller, local distances.

## Notes

### Competing Interest Statement

The authors have declared no competing interest.

